# PACT suppresses PKR activation through dsRNA binding and dimerization, and is a therapeutic target for triple-negative breast cancer

**DOI:** 10.1101/2025.05.27.656288

**Authors:** Addison A. Young, Isabelle G. Juhler, Jackson R. Pierce, Holly E. Bohlin, Haley A. Harper, David S. Onishile, Renee N. Chua, Madison E. Liu, Estelle N. Gardner, Bennett D. Elzey, Kyle A. Cottrell

## Abstract

Triple-negative breast cancer (TNBC), the deadliest breast cancer subtype, lacks broadly applicable targeted therapies. Induction of ‘viral mimicry’ by activation of viral double-stranded RNA (dsRNA) sensors has potential therapeutic applications for TNBC and other cancers. Suppressors of dsRNA sensing prevent sensing of endogenous dsRNAs and resulting autoimmunity. Depletion of the suppressor of dsRNA sensing ADAR1 causes activation of dsRNA sensors and cell death in many cancer cell lines. These ADAR1-dependent cells are generally also dependent on the dsRNA-binding protein PACT, which is highly expressed and essential in many TNBC cell lines. While PACT is known as an activator of the dsRNA sensor PKR, overexpression of PACT had no effect on activation of PKR in multiple TNBC cell lines. Conversely, depletion of PACT in PACT-dependent cell lines caused robust activation of the dsRNA sensor PKR and cell death, in addition to induction of integrated stress response genes and NF-κB targets. These phenotypes were entirely dependent on PKR. Rescue experiments revealed that PACT dimerization and dsRNA binding is required to suppress PKR activation. While depletion of PACT alone in ADAR1/ PACT-independent cell lines had no effect on PKR activation, combined depletion of both PACT and ADAR1 in those cell lines caused robust PKR activation and cell death, supporting a partially redundant role for ADAR1 and PACT in suppression of dsRNA sensing. Taken together, these findings support a vital role for PACT in suppressing PKR activation and highlight the therapeutic potential of targeting PACT to treat TNBC.

## Introduction

Triple-negative breast cancer (TNBC) is the deadliest form of breast cancer, with high rates of recurrence and metastasis (Curigliano and Goldhirsch 2011; Bianchini et al. 2016). A major factor driving the poor outcomes of TNBC patients is the lack of broadly applicable targeted therapies for TNBC (Curigliano and Goldhirsch 2011; Bianchini et al. 2016; Waks and Winer 2019). Immunotherapies, such as immune checkpoint blockade (ICB), have shown some efficacy in TNBC, but many tumors are resistant (Morad et al. 2021). Often, tumors with increased inflammation, known as immunologically ‘hot’ (immune inflamed), are more sensitive to ICB, relative to ‘cold’ (immune excluded) tumors (Chen and Mellman 2017). Several recent studies have shown that increasing inflammation within tumors can overcome resistance to ICB (Ishizuka et al. 2019; Jiang et al. 2019; Guirguis et al. 2023; Huang et al. 2024; Young et al. 2024). A great example of this approach is targeting the RNA editor ADAR1 to overcome resistance to ICB in mouse tumor models (Ishizuka et al. 2019).

Adenosine Deaminase Acting on RNA (*ADAR*, which encodes ADAR1) has been identified as an essential gene in multiple cancer cell lines – including those derived from breast, lung and ovarian cancer (Gannon et al. 2018; Liu et al. 2019; Kung et al. 2021). ADAR1 deaminates adenosine to inosine in double-stranded RNAs (dsRNA) in a process known as A-to-I editing (Bass and Weintraub 1988; Bass 2024; Mendoza and Beal 2024). This function of ADAR1 is essential and prevents autoimmunity. Specifically, A-to-I editing by ADAR1 prevents activation of MDA5, a double-stranded RNA (dsRNA) sensor, by endogenous dsRNAs (Liddicoat et al. 2015; Pestal et al. 2015). There are multiple dsRNA sensors expressed in immune and non-immune cells that detect dsRNA arising from viral infections (Rehwinkel and Gack 2020; Chen and Hur 2022; Cottrell et al. 2024a). Because these proteins bind to dsRNA by recognizing the structure of the dsRNA, and generally lack any sequence specificity, dsRNA sensors can also be activated by endogenous dsRNAs (Cottrell et al. 2024a). Several dsRNA sensors, including MDA5 and RIG-I, activate the type-I interferon (IFN-I) pathway to promote an antiviral response (Rehwinkel and Gack 2020). The dsRNA sensor protein kinase RNA-activated (PKR) instead activates the integrated stress response (ISR) by phosphorylation of eIF2α and promotes inflammation through activation of NF-kB (Gal-Ben-Ari et al. 2018; Chukwurah et al. 2021).

Since dsRNA sensors can bind and be activated by endogenous dsRNAs, to prevent autoimmunity, their activation must be suppressed in the absence of a viral infection. Multiple proteins suppress activation of dsRNA sensors by endogenous dsRNAs. These ‘suppressors of dsRNA sensing’, function through several mechanisms. For instance, ADAR1 prevents activation of two different dsRNA sensors through distinct mechanisms. Whereas ADAR1 prevents activation of MDA5 through A-to-I editing of dsRNA, ADAR1 also prevents activation of PKR through competition for dsRNA binding (Liddicoat et al. 2015; Pestal et al. 2015; Chung et al. 2018; Hu et al. 2023). The activation of dsRNA sensors and their downstream pathways, triggered by the loss of suppressors of dsRNA sensing or other perturbations, is referred to as ‘viral mimicry’. This term reflects cell behavior similar to that elicited by viral infection, but instead the cells are responding to endogenous dsRNAs (Chen et al. 2021; Cottrell et al. 2024a). It is this viral mimicry phenotype that overcomes resistance to ICB in ADAR1 depleted tumors and serves as a strong justification for the identification of ADAR1 inhibitors to treat multiple cancers (Ishizuka et al. 2019).

While the effects of depleting ADAR1 in cancer cells has therapeutic value, not all cancer cells are dependent on ADAR1 for proliferation (Gannon et al. 2018; Liu et al. 2019; Kung et al. 2021). We and others have observed that depletion of ADAR1 causes cell death and/or reduced proliferation and activation of dsRNA sensors in only a subset of cancer cell lines. Roughly half of TNBC lines are dependent on ADAR1 expression, based on reduced gene dependency scores from DepMap (McFarland et al. 2018; Dempster et al. 2019; Broad 2024a). In these ADAR1-dependent cell lines, depletion of ADAR1 causes cell death and activation of multiple dsRNA sensing pathways (Gannon et al. 2018; Liu et al. 2019; Kung et al. 2021). Conversely, in ADAR1-independent cell lines, depletion of ADAR1 has no effect on cell viability and there is no activation of dsRNA sensors. One factor driving ADAR1-dependency is elevated expression of IFN stimulated genes (ISGs) in ADAR1-dependent cell lines. The chronic IFN-I signaling in these cells leads to elevated expression of dsRNA sensors (PKR, MDA5, OAS1-3) that are ISGs (Gannon et al. 2018; Liu et al. 2019; Kung et al. 2021). It has been proposed that the elevated expression of dsRNA sensors in these cells creates a poised state, where the cells are highly sensitive to loss of ADAR1 – or potentially the loss of other suppressors of dsRNA sensing.

ADAR1 is not the only suppressor of dsRNA sensing. For example, DHX9 and STAU1 prevent PKR activation through binding to dsRNAs (Elbarbary et al. 2013; Cottrell et al. 2024b). Like ADAR1, those proteins bind dsRNA through dsRNA binding domains (dsRBD). There are nineteen human proteins that contain dsRBDs, and many other dsRNA binding proteins (dsRBP) that bind via other domains, in particular helicase domains. Each of these proteins are potentially competing with PKR, or perhaps MDA5, for binding to endogenous dsRNAs.

Here we provide evidence that <u>p</u>rotein <u>act</u>ivator of the interferon-induced protein kinase (PACT) functions as a suppressor of dsRNA sensing in TNBC. We show that PACT specifically suppresses PKR activation through dimerization and dsRNA binding. In addition to PKR activation and cell death, depletion of PACT causes activation of the ISR and NF-kB pathways in PACT-dependent cell lines. In PACT-independent cell lines, our data support redundancy between PACT and ADAR1 in suppression of dsRNA sensing. Together, our findings support PACT as a therapeutic target for a subset of TNBC.

## Results

### PACT is highly expressed in TNBC and essential in many TNBC cell lines

Since ADAR1-dependent cell lines have elevated expression of PKR and other dsRNA sensors, we hypothesized that they would be sensitive to loss of other suppressors of dsRNA sensing. As such, we turned to publicly available gene dependency data from DepMap to identify dsRBPs that may function as suppressors of dsRNA sensing like ADAR1. We determined the pairwise correlation coefficients for ADAR1-dependency scores vs the dependency scores for all other genes across all cell lines for which dependency data is available in DepMap (Broad 2024b). This analysis was performed for dependency scores from CRISPR-Cas9 screens (CHRONOS) and RNAi screens (DEMETER2) (McFarland et al. 2018; Dempster et al. 2019). Among the genes with the strongest correlation with ADAR1-dependency scores was *PRKRA*, which encodes PACT, (Pearson r = 0.341 for CHRONOS and 0.420 for DEMETER2), (Fig. 1a-b; Supplemental Fig. S1a-b). While there are many other genes with dependency scores that significantly correlate with ADAR1-dependency, including the RNA exonuclease XRN1 which has previously been shown to suppress activation of dsRNA sensors (Zou et al. 2024), we decided to focus our research on PACT because it, like ADAR1, contains multiple dsRNA binding domains. Additionally, PACT was the only dsRBD containing protein with gene dependency scores that strongly correlate (r > 0.2) with ADAR1-depedendency (Supplemental Fig. S1c).

**Figure 1:**
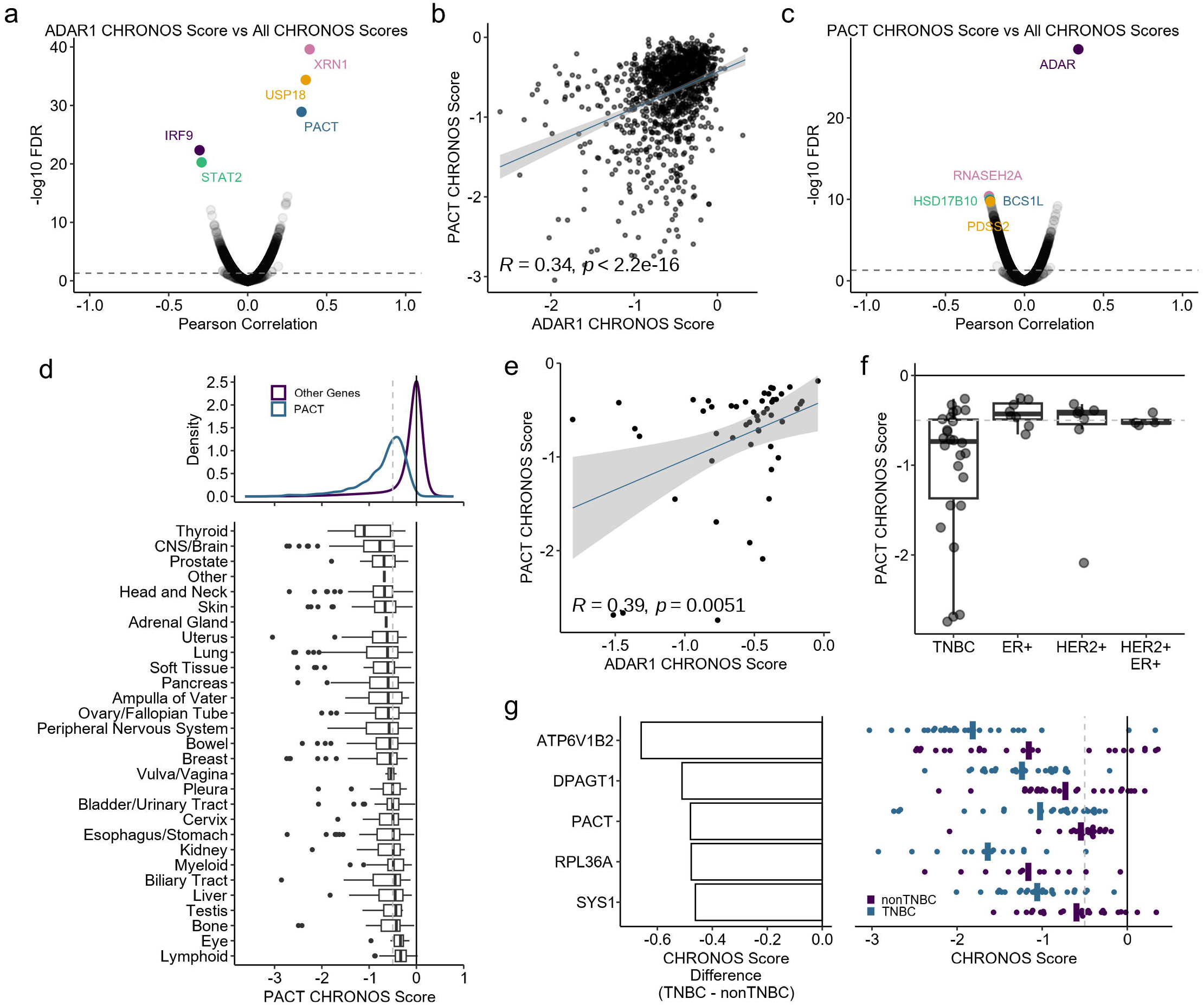
PACT is a co-dependency of ADAR1 and is essential in many TNBC cell lines. **a** and **c** Volcano plots of Pearson correlation coefficients and FDR corrected p-values for pairwise comparisons between ADAR1 CHRONOS score (**a**) or PACT CHRONOS score (**c**) and CHRONOS scores for all genes in DepMap across all cell lines. **b** Correlation between PACT and ADAR1 CHRONOS scores for all DepMap cell lines, Pearson correlation coefficient and p-value shown. **d** Top, density plot of CHRONOS scores for PACT or all other genes. Bottom, boxplots for PACT CHRONOS score by lineage. **e** Correlation between PACT and ADAR1 CHRONOS scores for breast cancer cell lines, Pearson correlation coefficient and p-value shown. **f** Boxplot of PACT CHRONOS Scores of breast cancer cell lines separated by subtype. **g** Left, CHRONOS score difference between TNBC and non-TNBC cell lines. Right, CHRONOS Score box plots for TNBC and non-TNBC cell lines. The top five genes based on the difference of CHRONOS scores between TNBC and non-TNBC are shown. All data shown from DepMap.

Analysis of pairwise correlation between PACT-dependency scores and those of all other genes revealed ADAR1 as the strongest co-dependent gene of PACT (Fig 1c; Supplemental Fig. S1d). Looking across lineages, PACT-dependent cell lines are common in cancers arising from multiple organs (Fig. 1d; Supplemental Fig. S1e). Given our past efforts studying ADAR1 in breast cancer, the strong correlation between PACT-dependency and ADAR1-depdency scores in breast cancer cell lines (Fig. 1e and Supplemental Fig. S1f), and several breast cancer cell lines being among the most strongly dependent on PACT, we chose to focus on studying the role of PACT in breast cancer. Analysis of PACT-dependency in breast cancer subtypes revealed a strong bias towards triple-negative breast cancer (TNBC) (Fig. 1f; Supplemental Fig. S1g-i). Interestingly, the difference between PACT-dependency scores amongst TNBC versus non-TNBC cell lines is larger than the difference for all but two other genes (Fig. 1g).

The expression of PACT at the RNA and protein levels varies across cell lines, with TNBC cell lines generally expressing more PACT than non-TNBC lines (Fig. 2a, Supplemental Fig. S2a-b). In human tumors, PACT expression is significantly elevated in TNBC relative to normal breast or other subtypes, (Fig. 2d-e and Supplemental Fig. S2c-d). Elevated expression of PACT, however, is not prognostic of survival in breast cancer overall, or TNBC (Fig. 2f-g).

**Figure 2:**
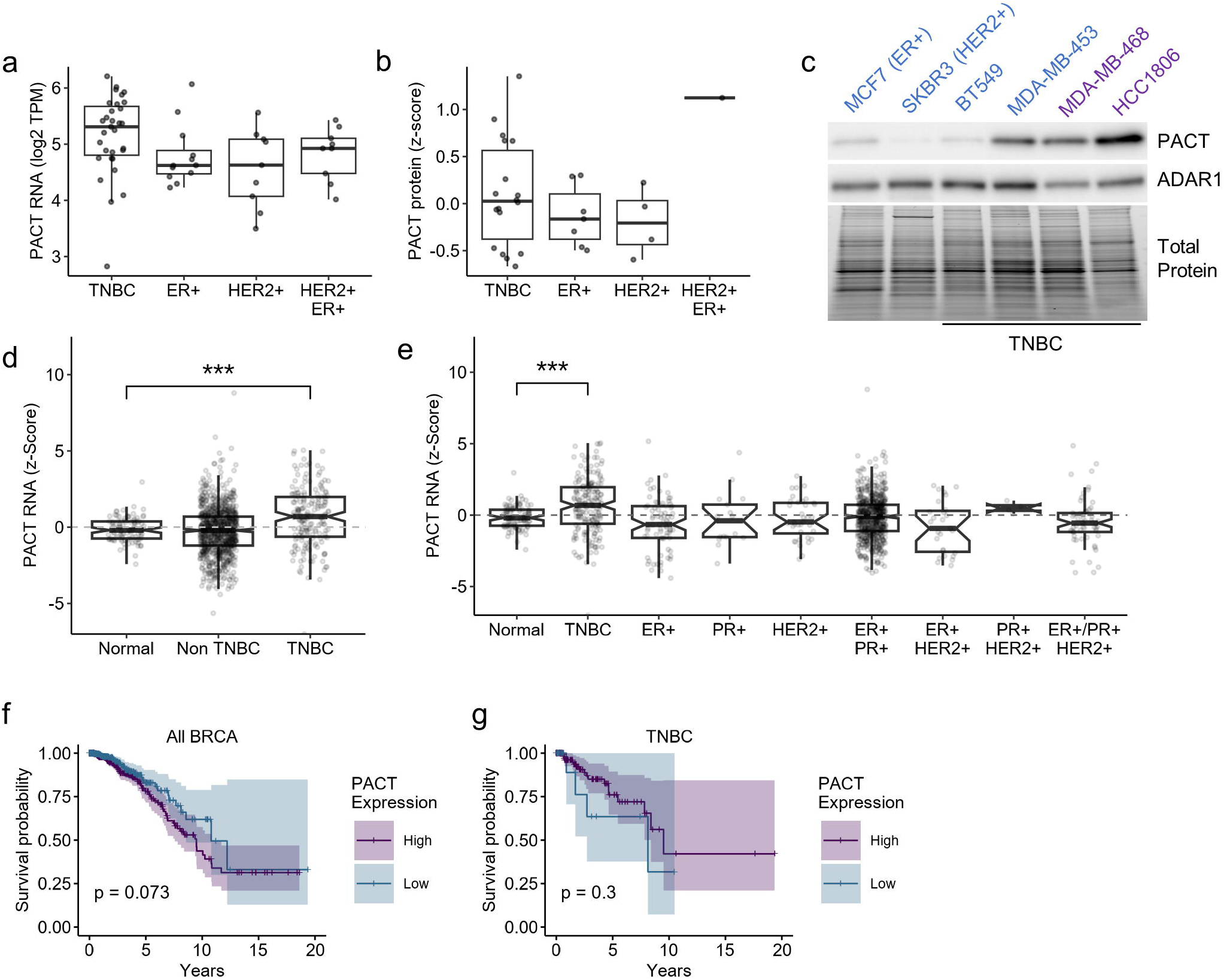
PACT is highly expressed in TNBC. Expression of PACT at the RNA (**a**) or protein level (**b**) in breast cancer cell lines separated by subtype. **c** Representative immunoblot for PACT and ADAR1 expression in a panel of breast cancer cell lines. Cell lines in blue are PACT-independent and purple are PACT-dependent. Total protein was imaged using a Stain-Free Gel and serves as a loading control. **d** – **e** Expression of PACT at the RNA level in normal and breast tumor samples separated by subtypes. **e** – **f** Overall survival of breast cancer patients separated by high or low PACT expression for all tumor types (**e**) or TNBC only (**f**). For panels **a**,**b**,**d** and **e**, data from DepMap; **f**, **g** from TCGA.

### PACT is not an activator of PKR in TNBC

Previous research has indicated that PACT is an activator of the dsRNA sensor PKR (Patel and Sen 1998; Patel et al. 2000; Peters et al. 2001; Li et al. 2006; Peters et al. 2009; Singh et al. 2011; Chukwurah et al. 2021). To evaluate if PACT functions as an activator of PKR in breast cancer cell lines, we overexpressed either wildtype PACT, or two phospho-site mutants of PACT (phosphomimetic S287D and phospho-null S287A). Those mutants alter a serine residue in PACT that was previously shown to enhance activation of PKR when phosphorylated (Peters et al. 2009; Singh et al. 2011). Overexpression of neither PACT, PACT^S287D^ nor PACT^S287A^ had any effect on PKR activation (phosphorylation of PKR on Thr- 446) in four different TNBC cell lines (Fig. 3a-b). To further evaluate if PACT functions as an activator of PKR in cancer, we compared PACT expression at the protein level with expression of ATF4, a component of the ISR that is induced upon PKR activation (Costa-Mattioli and Walter 2020). We observed no correlation between PACT and ATF4 protein expression across cancer cell lines for which proteomic data were available (Fig. 3c). As a control, we compared ATF4 expression to ATF3, a transcription factor induced by ATF4, and found a strong correlation (Fig. 3d). Together these findings do not support PACT functioning as a PKR activator.

**Figure 3:**
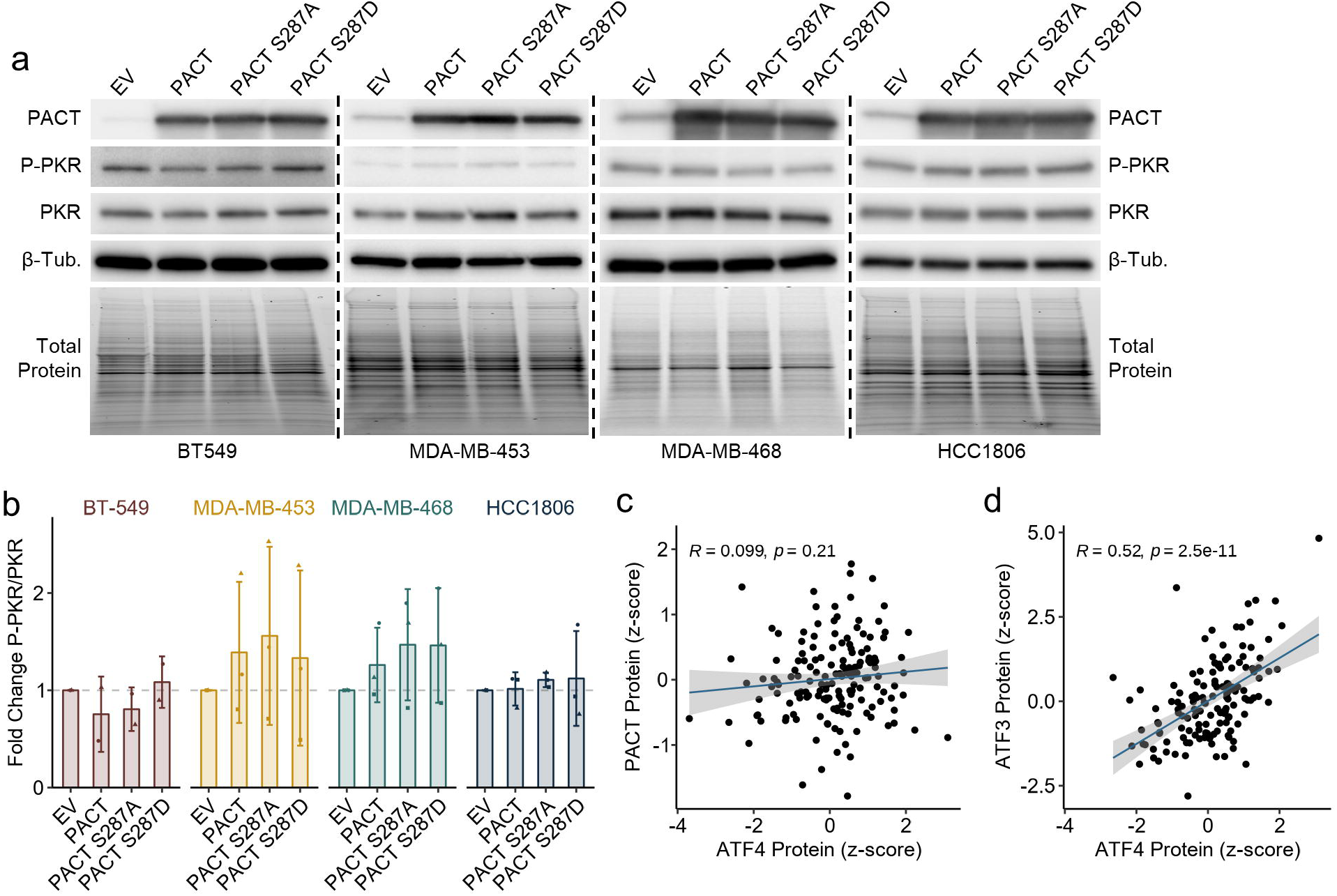
PACT overexpression does not cause PKR activation in TNBC. **a** Representative immunoblots for proteins of interest in control (empty vector, EV) or PACT (PACT, PACT S287A, PACT S287D) overexpressing cell lines. The blots for each cell line were performed independently and should not be compared between cell lines. Total protein was imaged using a Stain-Free Gel and was used as the loading control for normalization. **b** Quantification of the blot in **a**. Bars represent the average of at least three biological replicates, error bars are +/- SD. **c** and **d** Scatter plots comparing the protein abundance of PACT (**c**) or ATF3 (**d**) to ATF4 protein abundance across cancer cell lines, the Pearson correlation coefficient and p-value are shown. Data for **c** and **d** from DepMap.

### PACT suppresses PKR activation, but not activation of other dsRNA sensors

To further evaluate the role of PACT in TNBC cell lines, we used CRISPR-Cas9 to deplete PACT in a panel of PACT-dependent and PACT-independent cell lines (Fig. 4a-b). The PACT-dependent and -independent cell lines chosen are also ADAR1-dependent and - independent, respectively (Fig. 4a). Consistent with DepMap data, for PACT-dependent cell lines, depletion of PACT reduced cell viability, but in PACT-independent cell lines, depletion of PACT had no effect on viability as measured by an ATP dependent luciferase activity (Fig. 4c). For two PACT-dependent cell lines, we further evaluated the effect of PACT depletion on cell viability and proliferation by crystal violet staining. Depletion of PACT reduced crystal violet stained foci in both HCC1806 and MDA-MB-468 (Fig. 4d). To evaluate the *in vivo* effect of PACT depletion on tumorigenesis, we performed an orthotopic xenograft study with inducible knockout of PACT in HCC1806 cells. Tumorigenesis was significantly reduced by PACT depletion, highlighting the potential of targeting PACT to treat TNBC (Fig. 4e).

**Figure 4:**
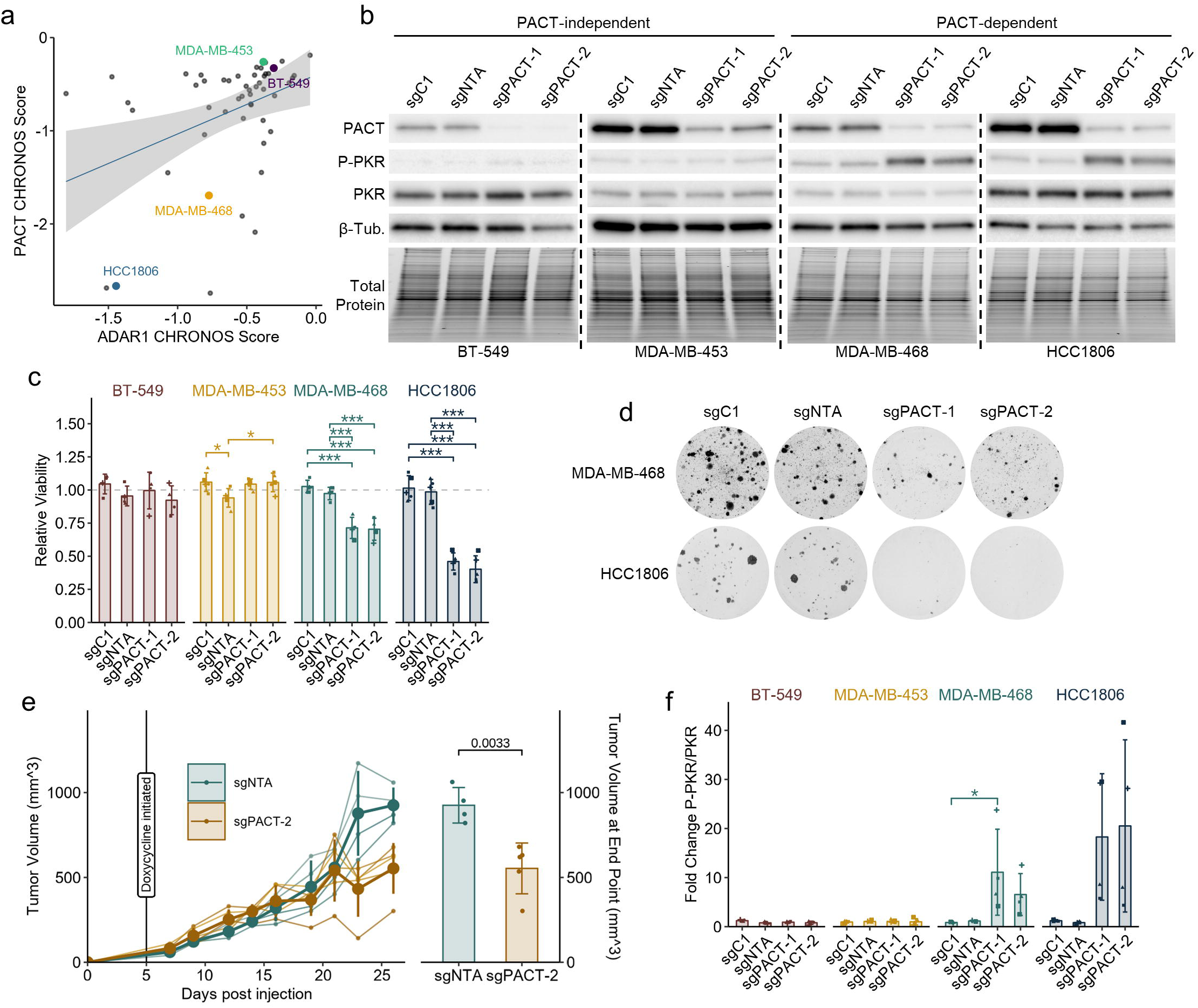
In PACT-dependent TNBC cells, PACT is required for viability, tumorigenesis, and suppression of PKR activation. **a** Correlation between PACT and ADAR1 CHRONOS scores of breast cancer cell lines, Pearson correlation coefficient and p-value shown, data from DepMap. The labeled cell lines are used in PACT depletion experiments. **b** Representative immunoblot for PACT-independent and PACT-dependent cell lines with (sgPACT-1, sgPACT-2) or without (sgNTA, sgC1) depletion of PACT. The blots for each cell line were performed independently and should not be compared between cell lines. Total protein was imaged using a Stain-Free Gel and was used as the loading control for normalization. **c** Cell viability as assessed by CellTiter-Glo 2.0 in PACT depleted and control cell lines. **d** Representative crystal violet staining of PACT depleted and control cells. **e** Effect of PACT depletion on tumorigenesis of HCC1806 cells. Left panel is tumor volume over time, right panel is the final tumor volume at end-point. **f** Quantificaiton of the immunoblot in **b**. Bars represent the average of at least four biological replicates, error bars are +/- SD. * p <0.05, ** p <0.01, *** p < 0.001. P-values determined by one-way ANOVA with post-hoc Tukey (**c** and **f**), or t-test (**e**).

In PACT-dependent cell lines, but not PACT-independent cell lines, depletion of PACT caused activation of PKR (Fig. 4b,f). This finding suggests that PACT functions not as an activator of PKR, but as a suppressor of PKR activation. We performed RNA-seq on PACT depleted and control HCC1806 and MDA-MB-468 cells to evaluate transcriptomic response to PACT depletion (Fig. 5a-b, Supplemental Tables 3-4). Activation of the dsRNA sensors MDA5, RIG-I and TLR3 leads to induction of type I IFN stimulated genes (ISG) (Matsumoto et al. 2011; Rehwinkel and Gack 2020). PACT depletion had little effect on type I ISG (Hallmark interferon alpha response (Liberzon et al. 2015)) expression in HCC1806 or MDA-MB-468 (Fig. 5c; Supplemental Fig. S3a-b). This observation was validated by qRT-PCR which again indicated little to no change in ISG expression in PACT depleted cells relative to controls, except for one ISG (IFIT2) which was elevated in the PACT-independent cell line MDA-MB-453 upon PACT depletion (Fig. 5d). The 2’-5’-oligodenylate synthases (OAS1, OAS2, and OAS3) are another group of dsRNA sensors that are also ISGs (Hovanessian and Justesen 2007). When activated, the OAS proteins activate RNase L which degrades many cellular RNAs, including rRNA leaving behind a distinctive banding pattern (Chakrabarti et al. 2011). Analysis of rRNA integrity in PACT-depleted and control cells revealed no cleavage that would be consistent with RNase L activation (Fig. 5e). Taken together, these findings suggest that in PACT-dependent cells, PACT suppresses activation of PKR, but not MDA5, RIG-I, TLR3 or the OASs.

**Figure 5:**
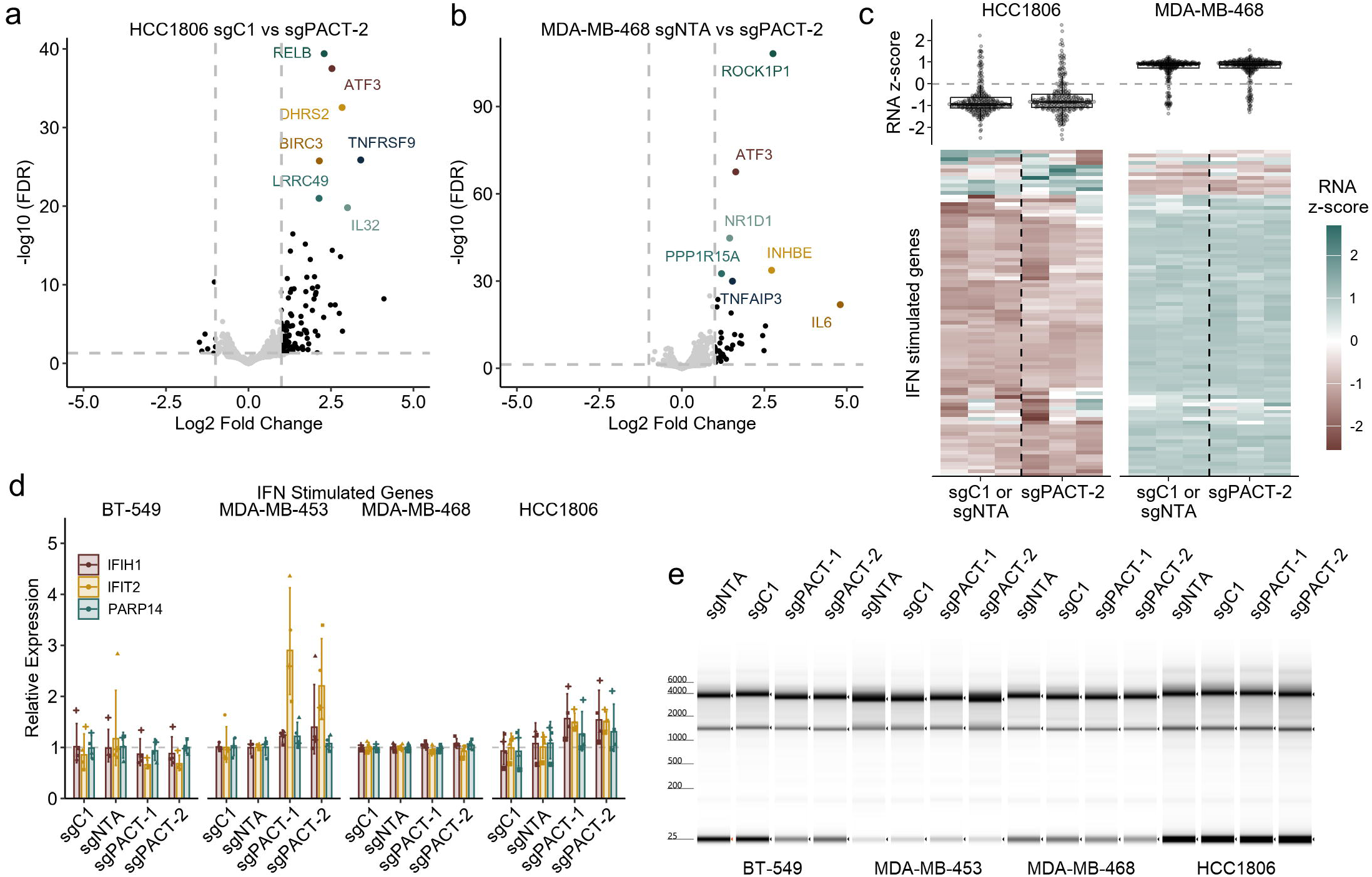
PACT depletion does not cause activation of type I IFN or RNaseL. **a** and **b** Volcano plot for fold-change of RNA expression in PACT depleted (sgPACT-2) or control (sgC1 or sgNTA) HCC1806 (**a**) or MDA-MB-468 (**b**). **c** Heatmap for RNA expression of type I ISGs (genes belonging to the gene set: Hallmark interferon alpha response). Top panel, box and overlayed ‘quasirandom’ plots for all ISGs in the heatmap below. The heatmap is clustered by gene (rows), the dendrogram has been omitted for brevity. **d** qRT-PCR for RNA expression for ISGs in PACT depleted and control cells. Bars represent the average of at least four biological replicates, error bars are +/- SD. **e** Representative psuedo-gel images of rRNA integrity for PACT depleted and control cells.

### Activation of PKR in PACT depleted cells drives cell death and activation of ISR and NF- κB

Gene set enrichment analysis of PACT depleted cells identified many dysregulated pathways. The top two most upregulated pathways in both HCC1806 and MDA-MB468 were associated with NF-κB signaling (Hallmark TNFA signaling via NFKB (Liberzon et al. 2015)) and the ISR (ATF4 target genes (Wong et al. 2019)). PKR is a well-known activator of the ISR, which is largely facilitated by the transcription factor ATF4 (Gal-Ben-Ari et al. 2018; Costa-Mattioli and Walter 2020). In PACT-depleted cells, we observed elevated expression of several ATF4 targets, consistent with activation of ISR, in both our RNA-seq data, (Fig. 6a; Supplemental Fig. S3c-d,g; Supplemental Tables 7-8) and by qRT-PCR (Fig. 6b). Immunoblot analysis confirmed upregulation of the ATF4 targets GADD34 (encoded by *PPP1R15A*) and ATF3 (Fig. 6d; Supplemental Fig. 3i). We did not observe robust phosphorylation of eIF2α as would be expected based on the ISR signature in our RNA-seq data and activation of PKR (Fig. 6d; Supplemental Fig. 3i). However, GADD34, an eIF2α phosphatase that is expressed during the ISR to dephosphorylate eIF2α to shut down the ISR (Novoa et al. 2001) was highly expressed in our cells at the time of harvest which could explain the lack of elevated p-eIF2α in PACT depleted cells (Fig. 6d).

**Figure 6:**
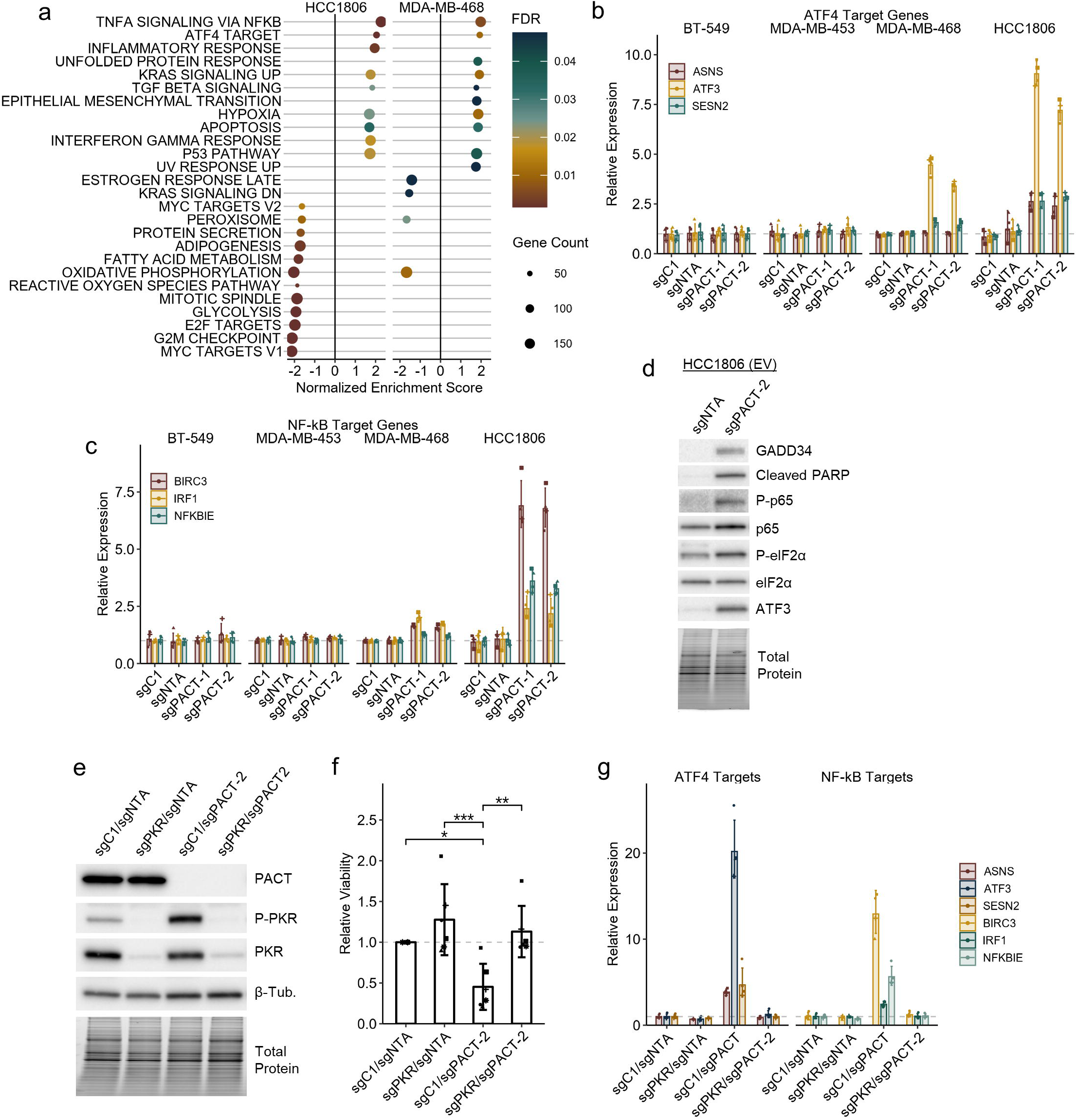
PKR is required for activation of NF-κB and ISR upon PACT depletion. **a** Dot blot summarizing the gene sets that are significantly up or downregulated in PACT depleted cells. FDR is the FDR corrected p-value for each gene set. **b** and **c,** qRT-PCR for RNA expression for ATF4 targets (**b**) or NF-κB targets (**c**) in PACT depleted and control cells. **d** Representative immunoblot of PACT depleted and control HCC1806 EV (empty vector) cells. Total protein was imaged using a Stain-Free Gel and was used as the loading control for normalization. **e** Representative immunoblot for control (sgC1/sgNTA), PACT depleted (sgC1/sgPACT-2), PKR depleted (sgC1/sgPKR) or combined depleted cell lines (sgPKR/sgPACT-2). **f** Cell viability as assessed by CellTiter-Glo 2.0 for the same conditions as **e**. **g** qRT-PCR for RNA expression for ATF4 targets or NF-κB targets for the same conditions as **e**. Bars represent the average of at least four biological replicates, error bars are +/- SD. * p <0.05, ** p <0.01, *** p < 0.001. P-values determined by Dunnett’s test.

In addition to the ISR, PKR can also activate the transcription factor NF-κB (Kumar et al. 1994; Gil et al. 1999; Bonnet et al. 2000; Zamanian-Daryoush et al. 2000; Gil et al. 2001; Bonnet et al. 2006; Chukwurah et al. 2021). Multiple NF-κB targets were upregulated upon PACT depletion in PACT-dependent cell lines as assessed by RNA-seq (Fig. 6a; Supplemental Fig. S3e-f,h; Supplemental Tables 7-8), and qRT-PCR (Fig. 6c). Immunoblot confirmed phosphorylation of NF-κB p65 (Ser468) in PACT depleted HCC1806 cells (Fig. 6d). Given that PKR activation can drive cell death, we performed a combined depletion experiment to determine if PKR activation is required for cell death upon PACT depletion (Fig. 6e). Depletion of PKR by CRISPR-Cas9 completely rescued the reduced viability caused by depletion of PACT in the PACT-dependent cell line HCC1806 (Fig. 6f). Furthermore, in PKR depleted cells, depletion of PACT did not cause activation of ATF4 or NF-κB target genes (Fig. 6g), indicating that elevated ATF4 and NF-κB target gene expression in PACT depleted cells is entirely dependent on PKR activation.

### PACT requires dimerization and dsRNA binding to suppress PKR activation

Multiple proteins suppress activation of PKR through binding to endogenous dsRNAs (Park et al. 1994; Elbarbary et al. 2013; Hu et al. 2023; Cottrell et al. 2024b). For instance, while ADAR1 prevents activation of MDA5 through A-to-I editing, it suppresses PKR activation through competition for dsRNA binding via its dsRBDs (Hu et al. 2023). To determine if dsRNA binding by PACT is required to suppress PKR activation, we performed a rescue experiment in which we overexpressed either wildtype PACT, or two dsRNA binding mutants of PACT. The mutants chosen, PACT-AA and PACT-EAA, were based on previous studies of PACT and ADAR1, in which two or three lysines of the KKxAK motif within their functional dsRBDs (the third dsRBD of PACT does not bind dsRNA (Peters et al. 2001)) had been mutated to abolish dsRNA binding (Valente and Nishikura 2007; Takahashi et al. 2013) (Fig. 7a). For both the wildtype and PACT mutants, we made synonymous mutations in the PACT coding sequence to prevent sgRNA binding. Depletion of PACT in empty vector (EV) control cells caused activation of PKR and reduced viability as before (Fig. 7b-d). Both phenotypes were completely rescued by overexpression of wildtype PACT, but neither of the PACT dsRNA binding mutants rescued, indicating that PACT suppresses PKR activation through binding endogenous dsRNAs (Fig. 7b-d).

**Figure 7:**
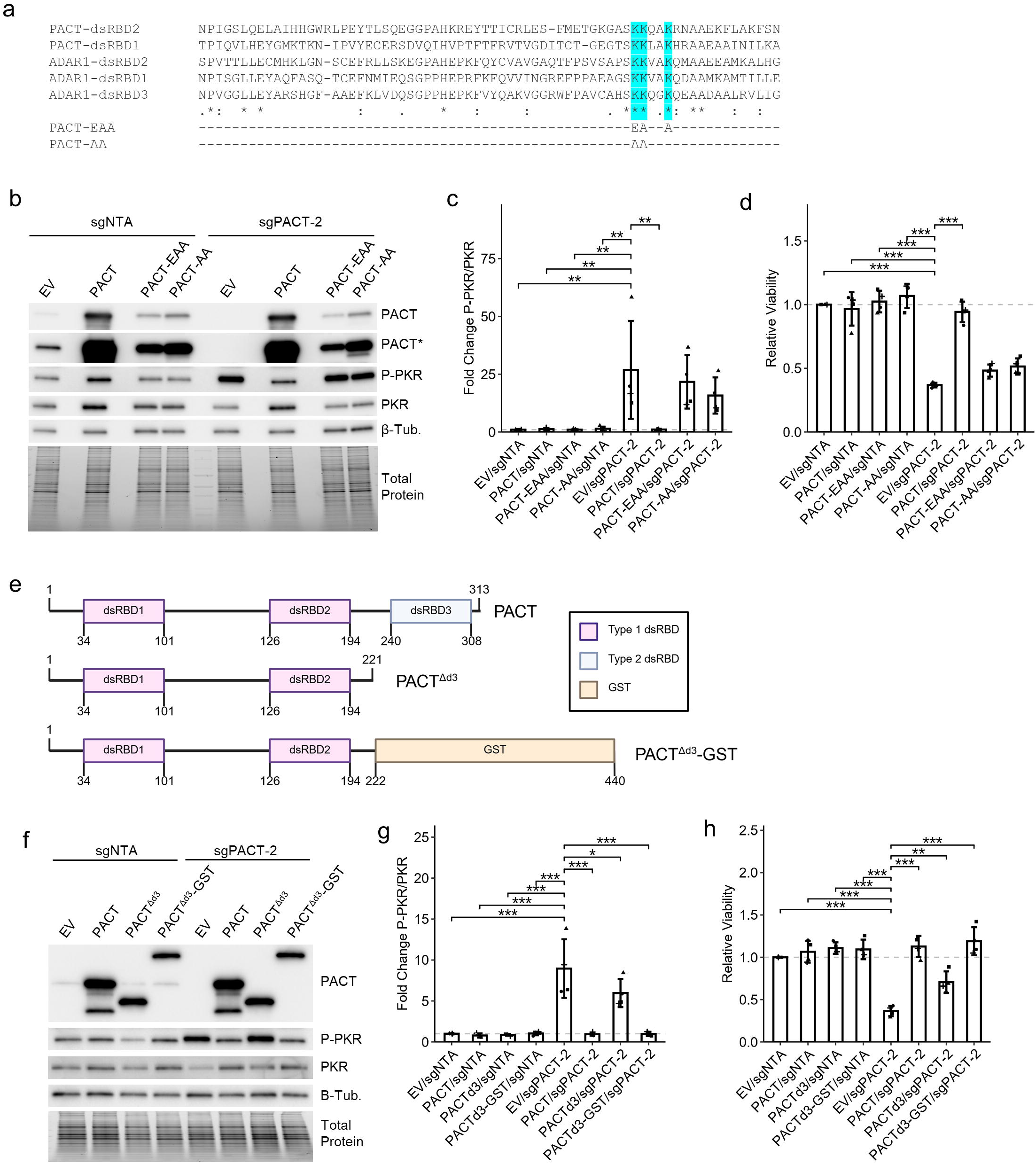
PACT dsRNA binding and dimerization is required for suppression of PKR activation. **a** Alignment of dsRBD 1 and 2 of PACT and dsRBDs 1-3 of ADAR1. The amino acid substitutions made for disrupting PACT dsRNA binding are shown below. **b** Representative immunoblot for control (sgNTA) and PACT depleted (sgPACT-2), with (PACT, PACT-EAA, PACT-AA) or without (EV, empty vector) overexpression of WT or dsRNA binding mutant PACT. **c** Quantification of the immunoblots in **b**. **d** Cell viability as assessed by CellTiter-Glo 2.0 for the same conditions as **b**. **e** Domain structure of WT PACT and PACT truncation and fusion constructs. **f** Representative immunoblot for control (sgNTA) and PACT depleted (sgPACT-2), with (PACT, PACT^Δd3^ or PACT^Δd3^-GST) or without (EV, empty vector) overexpression of WT or truncated PACT. **g** Quantification of the immunoblots in **f**. **h** Cell viability as assessed by CellTiter-Glo 2.0 for the same conditions as **f**. Bars represent the average of at least three replicates, error bars are +/- SD. * p <0.05, ** p <0.01, *** p < 0.001. P-values determined by Dunnett’s test.

Like PKR, PACT forms a homodimer, which is facilitated by its third dsRBD (Heyam et al. 2017). To evaluate if dimerization of PACT is required for its ability to suppress PKR activation, we performed another knockout-rescue experiment with two different PACT constructs. For the first construct, PACT^Δd3^, we truncated PACT to remove its third dsRBD, which is required for PACT dimerization (Heyam et al. 2017). To orthogonally restore dimerization of PACT^Δd3^, we generated a fusion construct that contained truncated PACT fused to GST to generate PACT^Δd3^-GST (Fig. 7e). As GST forms a strong dimer, this construct would be expected to restore dimerization of PACT^Δd3^. Like WT PACT, PACT^Δd3^ and PACT^Δd3^-GST maintained cytoplasmic localization (Supplemental Fig. 4a). When expressed in PACT depleted cells, PACT^Δd3^ did not rescue PKR activation or fully rescue cell viability (Fig. 7f-h). Conversely, PACT^Δd3^-GST completely rescued both PKR activation and cell viability (Fig. 7f-h), as well as ISR and NF-κB activation (Supplemental Fig. 4b). These findings indicate that while dsRBD3 is required for suppression of PKR activation, its function regarding suppression of PKR activation is only to allow for dimerization of PACT – which is required to suppress PKR activation.

### PACT and ADAR1 redundantly suppress PKR activation in PACT/ADAR1-independent cell lines

Given that PACT and ADAR1 are co-dependent genes in many cancer cell lines, and that they both rely on dsRNA binding to suppress PKR activation, we hypothesized that depletion of both ADAR1 and PACT in a PACT/ADAR1-independent cell line would cause activation of PKR. To evaluate that hypothesis, we depleted both PACT and ADAR1, each individually, or neither, in two PACT/ADAR1-independent TNBC cell lines, BT-549 and MDA-MB-453. As in our previous experiments, depletion of PACT alone in both cell lines had little to no effect on cell viability or PKR activation, and we observed the same for depletion of ADAR1 (Fig. 8a-c). Conversely, combined depletion of both ADAR1 and PACT in both cell lines caused robust activation of PKR and for BT-549 reduced viability. Consistent with PKR activation, we observed increased expression of ATF4 targets only in the PACT and ADAR1 depleted cells (Fig. 8d). Unlike in PACT-dependent cell lines, we did not observe substantial changes in NF-kB target expression (Fig. 8e). While the expression of most ISGs evaluated remained unchanged upon depletion of PACT and/or ADAR1 in each cell line, the ISG IFIT2 was upregulated upon PACT depletion in MDA-MB-453 and was further induced in the combined depleted cells (Fig. 8f). Finally, we observed no rRNA degradation consistent with OAS/RNase L activation upon depletion of PACT and/or ADAR1 in either cell line (Fig. 8g).

**Figure 8:**
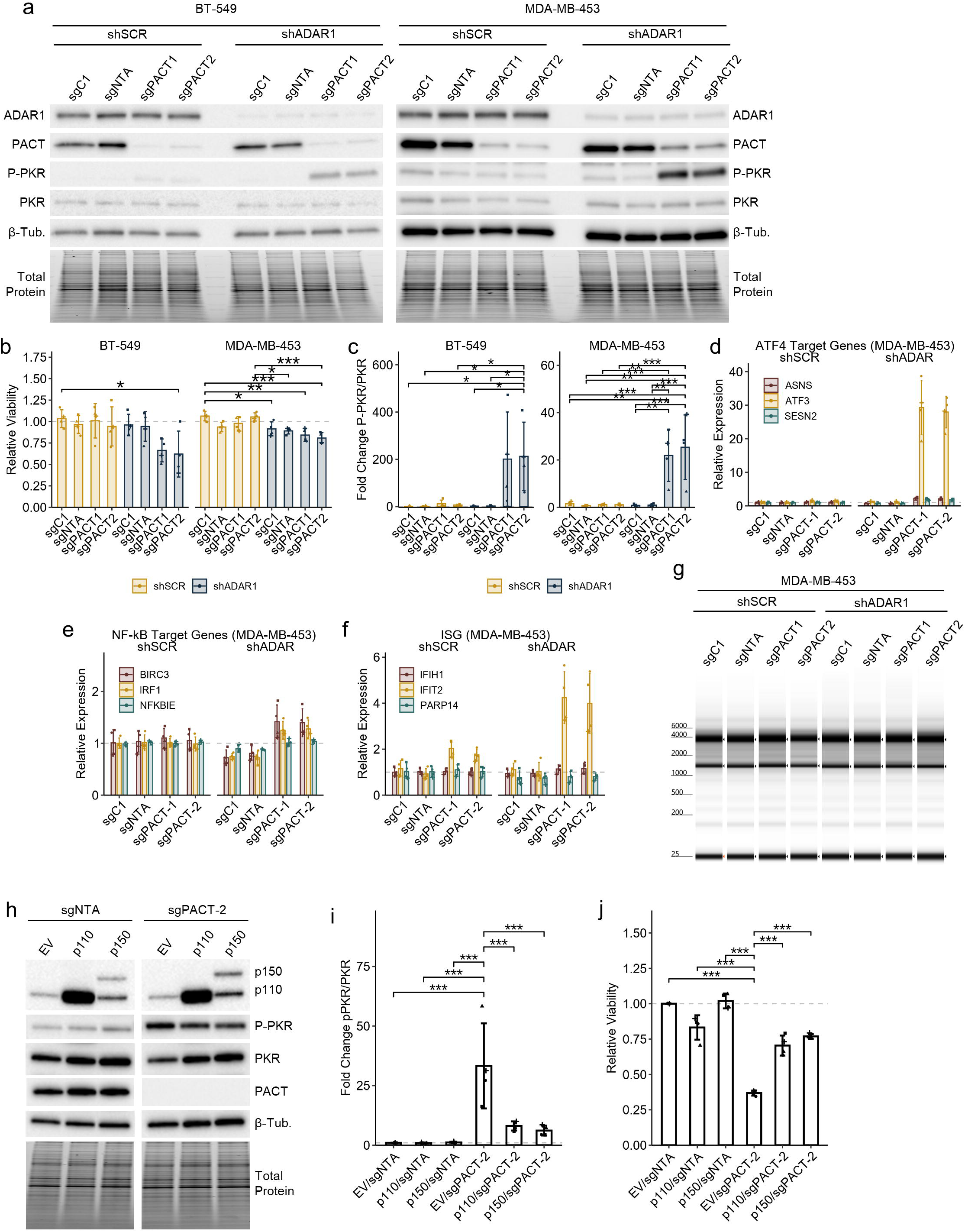
PACT and ADAR1 function redundantly to suppress PKR activation in PACT/ADAR1-independent cell lines. **a** Representative immunoblot for control (sgC1, sgNTA) and PACT depleted (sgPACT-1, sgPACT-2), with (shADAR1) or without (shSCR) knockdown of ADAR1. **b** Cell viability as assessed by CellTiter-Glo 2.0 for the same conditions as **b**. **c** Quantification of the immunoblots in **a**. **d**-**f** qRT-PCR for RNA expression of ATF4 targets (**d**), NF-κB targets (**e**) or ISGs (**f**) for the same conditions as **a**, MDA-MB-453. **g** Representative psuedo-gel images of rRNA integrity for the same conditions as **a**, MDA-MB-453. **h** Representative immunoblot for control (sgNTA) and PACT depleted (sgPACT-2), with (p110, p150) or without (EV, empty vector) overexpression of ADAR1 isoforms. **i** Quantification of the immunoblots in **h**. **j** Cell viability as assessed by CellTiter-Glo 2.0 for the same conditions as **h**. Bars represent the average of at least three replicates, error bars are +/- SD. * p <0.05, ** p <0.01, *** p < 0.001. P-values determined by one-way ANOVA with post-hoc Tukey (**b** and **c)**, or Dunnett’s test (**i** and **j**).

To further evaluate redundancy between PACT and ADAR1, we attempted to rescue the effects of PACT depletion in PACT-dependent cells by overexpression of each isoform of ADAR1. Overexpression of either the p110 or p150 isoform of ADAR1 partially rescued PKR activation and reduced viability caused by depletion of PACT in HCC1806 cells (Fig. 8h-j). Taken together with the combined depletion experiments above, these data support a redundant role for PACT and ADAR1 in suppression of PKR activation in PACT/ADAR1- independent cell lines.

### PACT-dependency correlates with PKR expression, which is elevated in TNBC

While the data above provide strong evidence to support PACT as a suppressor of PKR activation, they do not explain the differential sensitivity of PACT-dependent and PACT- independent cell lines to depletion of PACT. For ADAR1-dependency, it has been proposed that chronic IFN signaling in ADAR1-depenendent cells, which drives elevated expression of the dsRNA sensors MDA5, RIG-I, PKR and the OASs, sensitizes those cells to depletion of ADAR1 (Liu et al. 2019). This model likely does not explain PACT-dependency, because unlike ADAR1- dependency, there is no correlation between PACT-dependency score and ISG expression (Fig. 9a-b; Supplemental Fig. S5a-b). By analyzing proteomics data for cancer cell lines, we found that PACT-dependency strongly correlates with PKR expression at the protein level (Fig. 9c and Supplemental Fig. S5c-g). The correlation between PACT-dependency score and PKR protein expression is strongest in breast cancer (Fig. 9d). Consistent with the proteomics and DepMap data, we observed higher PKR expression in PACT-dependent cells relative to PACT- independent cells (Fig. 9e). Based on mass-spectrometry data, PKR expression is highest in TNBC cell lines (Fig. 9f) and in human tumors, PKR expression at the RNA level is elevated in breast tumors relative to normal breast, with TNBC tumors generally having higher expression (Fig. 9g-h; Supplemental Fig. S5h-i). While not as predictive as PKR expression, it should be noted that the expression of PACT itself correlates with PACT-dependency score (Supplemental Fig. S5j-m).

**Figure 9:**
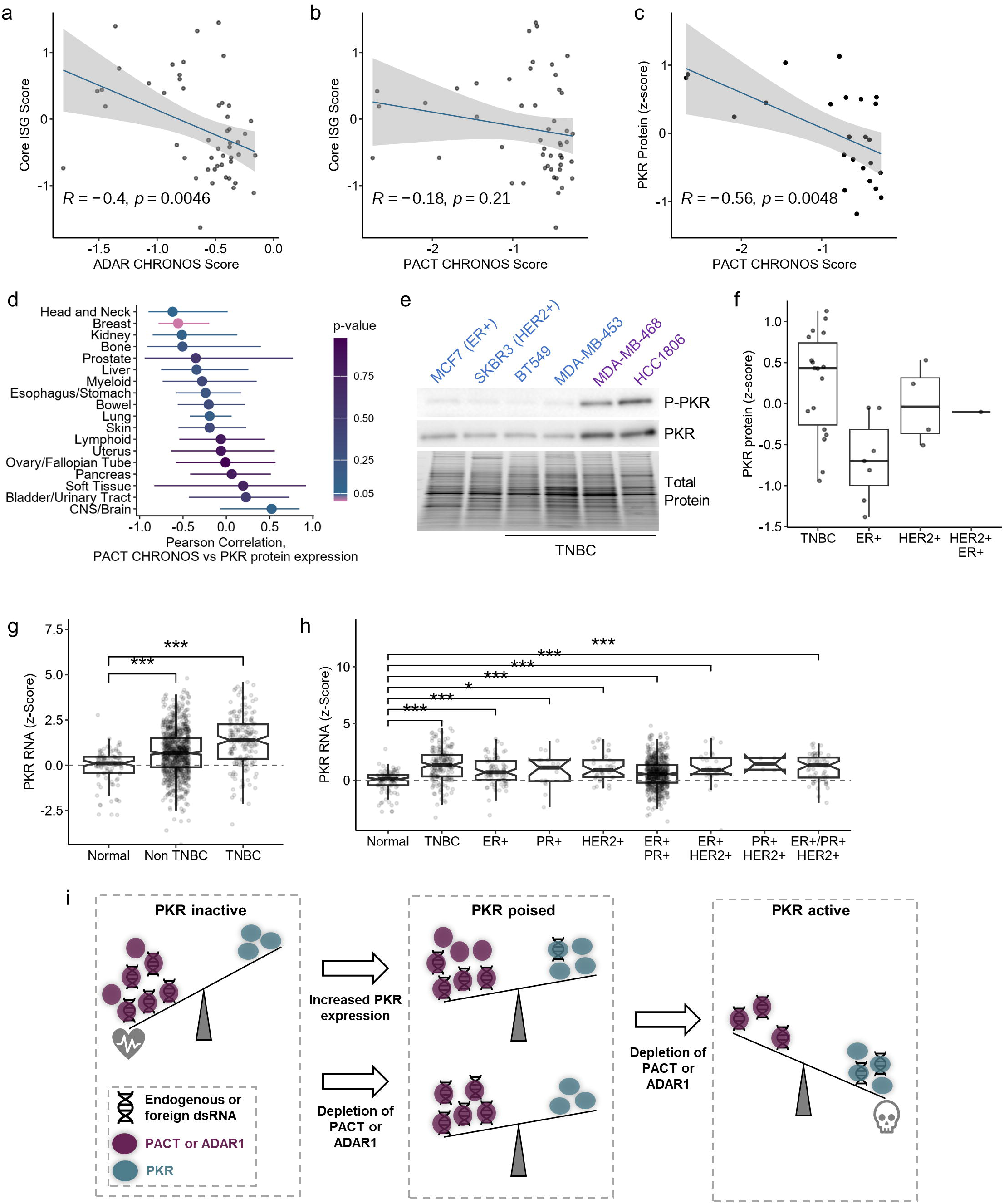
PKR expression is elevated in breast cancer and correlates with PACT-dependency. **a** – **b** Scatter plots comparing ISG expression (Core ISG Score, described previously (Kung et al. 2021)) and either ADAR1-dependency score (ADAR CHRONOS Score, **a**) or PACT-dependency score (PACT CHRONOS Score, **b**). **c** Scatter plot comparing PKR protein abundance and PACT-dependency score. For all scatter plots, the Pearson correlation coefficient and p-values are shown. **d** Summary of Pearson correlation coefficients and p-values between PKR protein abundance and PACT-dependency score for cancer cell line lineages. Lineages with PKR abundance and PACT-dependency scores for fewer than four cell lines were omitted. **e** Representative immunoblot for PKR and P-PKR in TNBC cell lines. Total protein is a Stain-Free gel image used as a loading control. **f** PKR protein abundance in breast cancer cell lines separated by subtype. **g** and **h** PKR RNA expression in normal human breast and breast tumors, data from TCGA. **i** Model for cellular sensitivity to depletion of ADAR1 and/or PACT. For panels **a**-**d** and **f**, data from DepMap; **g**-**h** from TCGA.

## Discussion

The role of PACT in regulation of PKR has been controversial in the literature. Early reports, which gave PACT its name, provided compelling evidence that PACT functioned to activate PKR through protein-protein interactions in the absence of dsRNA (Patel and Sen 1998; Peters et al. 2001; Li et al. 2006; Peters et al. 2009). Many groups have reported similar findings, often with PACT functioning to activate PKR during stress (Patel et al. 2000; Singh et al. 2011; Farabaugh et al. 2020; Chukwurah et al. 2021). However, other studies indicated that PACT conducts the exact opposite function, inhibiting PKR, including a mouse study showing that PKR (*Eif2ak2*) knockout could rescue embryonic lethality of PACT (Rax in mice, *Prkra*) knockout (Clerzius et al. 2013; Dickerman et al. 2015; Meyer et al. 2018). During the preparation of this manuscript, two studies were published that provided more definitive evidence, consistent with our data here, in support of PACT’s role as a suppressor of PKR activation (Ahmad et al. 2025; Manjunath et al. 2025).

While our data indicate that PACT suppresses PKR activation through dimerization and dsRNA binding, the finer details of the mechanism are not clear from our work alone. Fortunately, another group has examined the mechanism and reported those findings during the preparation of this manuscript (Ahmad et al. 2025). In this work, Ahmad, Zou, and Xhao et al., reveal that PACT suppresses PKR activation by preventing dimerization of sliding PKR monomers. Specifically, they propose a model in which weak protein-protein interactions between PKR and PACT prevent sliding of PKR along dsRNA thus preventing monomers from colliding and forming an active PKR dimer. While that model is supported by our own data showing that PACT requires dsRNA binding to prevent PKR activation, it is not clear how dimerization of PACT fits into the model. A recent study, published during the preparation of this manuscript, found that dimerization of the zebrafish homologue of PACT (Prkra) facilitated binding to and inhibition of eIF2 (Lu et al. 2025). In the model presented with those findings, Prkra uses its third dsRBD to dimerize and sequester eiF2 to inhibit translation, thus assigning a dual role for dsRBD3. Here, we found that dsRBD3 of PACT can be functionally replaced by GST (a dimeric protein), indicating that at least in human cells, dsRBD3’s only function in relation to suppression of PKR activity is to facilitate dimerization of PACT. How dimerization enables PKR inhibition is unclear. It is unlikely that dimerization allows PACT to better ‘compete’ with PKR for binding dsRNA as PACT mutants lacking the third dsRBD bind a model dsRNA with the same affinity as full-length PACT (Ahmad et al. 2025). Structural studies are needed to understand the role of dimerization; perhaps dimerization exposes the PKR binding site on PACT, or allows for higher-order structures of PACT-PKR-dsRNA that inhibit or sequester PKR.

Based on the elevated expression of PKR in PACT-dependent cells, we propose a model in which PACT-dependent cells are in a poised state, where because of the elevated abundance of PKR, the cell is highly sensitive to depletion of PACT, and in many cases ADAR1 (Fig. 9i). Conversely, in PACT-independent cell lines, the lower expression of PKR enables either PACT or ADAR1 alone to suppress PKR activation. In these cells, PACT and ADAR1 are functioning redundantly. This shared role of PACT and ADAR1 was also reported in a recent study published during preparation of this manuscript (Manjunath et al. 2025). This redundancy between ADAR1 and PACT is not unique. We have observed the same paradigm before for DHX9 and ADAR1, where depletion of DHX9 in ADAR1-dependent cells causes activation of PKR, while in ADAR1-independent cells, DHX9 and ADAR1 function redundantly to suppress activation of PKR and other dsRNA sensors (Cottrell et al. 2024b). A common thread connecting these proteins is the presence of dsRBDs. Here we show that the dsRNA binding activity of PACT is required for suppression of PKR activation, the same has been observed for ADAR1 previously (Hu et al. 2023), and for DHX9 expression of its dsRBDs alone is sufficient to suppress PKR activation (Cottrell et al. 2024b). While previous cellular studies indicate that ADAR1 suppresses PKR activation by competing with PKR for dsRNA binding, but the same was not observed *in vitro* by Ahmad, Zou, and Xhao et al. (Hu et al. 2023; Ahmad et al. 2025). Further studies are needed to evaluate in finer detail the mechanism by which ADAR1 suppresses PKR activation, and how that may or may not differ from PACT. More research is also needed to understand the role of ADAR1-p110 in suppressing PKR activation. Previous literature supports p150 as the ADAR1 isoform that suppresses PKR activation (Chung et al. 2018; Hu et al. 2023), though here we show that overexpression of p110 in PACT depleted cells reduces PKR activation. We have observed something similar in DHX9 and ADAR1 depleted cells, where expression of p110 was capable of suppressing PKR activation (Cottrell et al. 2024b). Additionally, Sinigaglia et al. observed reduced PKR activation upon p110 overexpression in A549 cells (Sinigaglia et al. 2024). In that study, a different model for inhibition of PKR by ADAR1 is proposed, a model in which the third dsRBD of ADAR1 binds the kinase domain of PKR preventing its activation. It is of course possible that suppression of PKR by p110 in these contexts is an artifact of overexpression and does not occur physiologically. Or perhaps, p110 utilizes this function in specific contexts when p150 or other proteins are unavailable, possibly in the nucleus (to which some PKR localizes, Supplemental Fig. S4a) (Jeffrey et al. 1995) or during mitosis, during which nuclear RNAs can activate PKR (Kim et al. 2014).

Finally, here we provide strong evidence to support the therapeutic targeting of PACT to treat TNBC. There are several attributes that make PACT a good target for TNBC, 1) PACT is highly expressed in TNBC, 2) PACT is essential in many TNBC cell lines, 3) PKR expression, which correlates with PACT-dependency, is elevated in TNBC. While PACT has no known enzymatic functions, disrupting PACT dimerization would inhibit its regulation of PKR. As such, PACT dimerization inhibitors, or PACT degraders, could have therapeutic potential for TNBC, and likely many other types of cancer.

## Materials and Methods

### Cell culture

Cell lines (MCF-7 (RRID:CVCL_0031), SK-BR-3 (RRID:CVCL_0033), BT-549 (RRID: CVCL_1092), HCC1806 (RRID: CVCL_1258), MDA-MB-468 (RRID: CVCL_0063) MDA-MB-453 (RRID: CVCL _0418), and 293T (RRID: CVCL_0063) were obtained from American Type Culture Collection which used STR profiling to authenticate the cell lines, all cell lines were obtained between 2023 and 2024. The cell lines 293T and SK-BR-3 were cultured in Dulbecco’s modified Eagle’s medium (DMEM) (Hyclone) with 10% fetal bovine serum (BioTechne), 2 mM glutamine (Hyclone), 0.1 mM nonessential amino acids (Hyclone), and 1 mM sodium pyruvate (Hyclone). The cell lines MDA-MB-453 and MDA-MB-468 were cultured in Leibowitz L-15 media (HyClone Cat# SH30525) with 10% fetal bovine serum (BioTechne). The cell lines HCC1806 and BT-549 were cultured in Roswell Park Memorial Institute 1640 media (Corning Cat# 10-041-CV) with 10% fetal bovine serum (BioTechne), for BT-549 recombinant insulin (Gibco) was added to 0.78 μg/mL. The cell line MCF-7 was cultured in modified Eagle’s media (HyClone Cat# SH30024.01) with 10% fetal bovine serum (BioTechne) and recombinant insulin at 10.2 µg/mL. Except for MDA-MB-453 and MDA-MB-468, all cell lines were grown at 37 °C at 5% CO_2_, MDA-MB-453 and MDA-MB-468 were grown at 37 °C with atmospheric CO_2_. Mycoplasma testing was performed by a PCR based method. All experiments were performed with cells under twenty passages.

### Viral Production and Transduction

Lentivirus was produced by LipoFexin (Lamda Biotech) or Polyethylenimine (branched, ∼25,000 Da, Sigma-Aldrich) transfection of 293T cells with pCMV-VSV-G (a gift from Bob Weinberg (Stewart et al. 2003), Addgene plasmid #8454; RRID:Addgene_8454) or pMD2.G (a gift from Didier Trono, Addgene plasmid #12259; RRID:Addgene_12259) and pSPAX2 (a gift from Didier Trono, Addgene plasmid #12260; RRID:Addgene_12260), and a transfer plasmid for expression of genes of interest, shRNAs, or sgRNAs. Culture media was changed the day after transfection and lentivirus containing media was collected the following day, or after two days. Lentivirus containing media was filtered through a 0.45 µm filter before transduction of cells of interest in the presence of 10 µg/mL protamine sulfate (Sigma-Aldrich). Depending on the transfer plasmid used and cell line, cells were selected with puromycin at 2 µg/mL (Sigma-Aldrich), 150 μg/mL hygromycin (Gibco or InvivoGen) 10 µg/mL Blasticidin (Fisher or InvivoGen), or 500 µg/mL G418 (InvivoGen).

### Plasmids

For all sgRNAs targeting genes of interest, oligos encoding the sgRNAs (Supplemental Table 1) were cloned into lentiGuide-puro (a gift from Feng Zhang ^44^, Addgene #52961; RRID:Addgene_52963), lenti-sgRNA-hygro (a gift from Brett Stringer (Stringer et al. 2019), Addgene #104991; RRID:Addgene_104991), or tet-pLKO-sgRNA-puro (a gift from Nathanael Gray (Huang et al. 2017), Addgene plasmid # 104321; RRID:Addgene_104321) by ligation of annealed and phosphorylated oligos into a restriction enzyme digested and dephosphorylated vector. The sgRNA sequence for lenti-sgPKR-hygro was used previously (Zou et al. 2024). The pLKO-shSCR-hygro and pLKO-shADAR1-hygro plasmids have been described previously (Cottrell et al. 2024b).

Two control sgRNAs were used in this study. One (sgC1) targets a control genomic locus (AAVS1) and has been used previously (Zou et al. 2024). The second control sgRNA (sgNTA) is a non-targeting sgRNA described previously (Doench et al. 2016). For sgNTA, lentiGuide-sgNTA-puro was purchased from Addgene (a gift from John Doench & David Root (Doench et al. 2016), Addgene plasmid #80248; RRID:Addgene_80248). All other plasmids for the control sgRNAs were cloned in the same manner as described above, and all sequences are available in Supplementary Table 1.

The PACT coding sequence flanked by BamHI and MluI restriction enzyme sites, a Kozak sequence (5’-CACC-3’), and containing wobble mutations to prevent targeting of sgRNAs was synthesized by TwistBio. Restriction enzyme digest and ligation was used to clone PACT into pLV-EF1a-IRES-Blast vector (a gift from Tobias Meyer ^45^, Addgene plasmid #85133; RRID:Addgene_85133). The PACT dsRBD mutants were cloned in the same manner, all coding sequences can be found in the Supplemental Information. The PACT-dsRBD3 constructs were made through restriction enzyme digest and ligation. For the PACT^Δd3^ plasmid, pLV-EF1a-PACT-IRES-Blast was digested with EcoRI, which cuts between the coding sequence for dsRBD2 and dsRBD3 within PACT, and 3’ of PACT within pLV-EF1a-IRES-Blast. A short double-stranded DNA oligonucleotide encoding two stop codons was ligated in frame with the PACT coding sequence to generate pLV-EF1a-PACT^Δd3^-IRES-Blast. The plasmid for PACT^Δd3^-GST was cloned in the same manner with GST in place of the oligonucleotide insert. GST was PCR amplified from pDEST15, kindly provided by Dr. Mark Hall, Purdue University. The PCR primers and oligonucleotides used for cloning the PACT-dsRBD3 constructs, as well as the coding sequences of the final constructs can be found in Supplementary Table 1 and the Supplemental Information. The ADAR1 overexpression constructs used here (pLV-EF1-Blast-p110 and pLV-EF1-Blast-p150) were cloned previously (Cottrell et al. 2024b).

All plasmids were confirmed by restriction enzyme digest, as well as Sanger sequencing and Nanopore whole plasmid sequencing.

### Genetic Depletion by CRISPR-Cas9

For all CRISPR-Cas9 depletion experiments, the cell lines used in this study were transduced with lentivirus for inducible expression of Cas9 (iCas9) using the transfer plasmid lenti-iCas9-neo (a gift from Qin Yan (Cao et al. 2016), Addgene plasmid # 85400; RRID:Addgene_85400). After transduction and selection with G418, cells with high GFP expression upon doxycycline induction were sorted on a BD FACS Aria by the Purdue Flow Cytometry and Cell Separation Facility. We used either constitutively expressed or inducible sgRNAs targeting genes of interest. For constitutively expressed sgRNAs, we observed premature knockout of the genes of interest in uninduced cells, likely due to ‘leaky’ expression of Cas9. As such, for all experiments utilizing constitutive sgRNA expression (single knockout of PACT in Fig. 4, or combined knockout of PACT and knockdown of ADAR1 in Fig. 8), we began each biological replicate by transducing cells with the lentivirus for sgRNA expression followed by selection and induction with doxycycline. The timelines for transduction, selection, induction, harvesting of cells and evaluation of cell viability for each cell line is described in Supplemental Table 2.

For all knockout-rescue experiments and combined knockout of PACT and PKR, we combined iCas9 with an inducible sgRNA construct (tet-pLKO-sgRNA-puro) which prevented knockout prior to induction (data not shown). PACT overexpression and empty vector (EV) control HCC1806-iCas9 cells were generated by lentiviral transduction and selection for transgene incorporation with blasticidin. Subsequently, PACT overexpression and control EV HCC1806-iCas9 lines were transduced with lentivirus made with tet-pLKO-sgPACT-2-puro or tet-pLKO-sgNTA-puro and selected with puromycin. Experimental replicates were initiated by doxycycline treatment and the timeline is described in Supplemental Table 2. For PKR knockout, HCC1806-iCas9 cells were first transduced with lentivirus made with lenti-sgC1-hygro, lenti-sgPKR-1-hygro or lenti-sgPKR-2-hygro and selected with hygromycin. Cas9 expression was induced by doxycycline and PKR knockout was confirmed by immunoblot (data not shown). The HCC1806-iCas9 control and PKR knockout lines were passaged without doxycycline before transduction with lentivirus made with tet-pLKO-sgPACT-2-puro or tet-pLKO-sgNTA-puro and selected with puromycin. Experimental replicates were initiated by doxycycline treatment and the timeline is described in Supplemental Table 2.

### Immunoblot

Cell pellets were lysed and sonicated in RIPA Buffer (50 mM Tris pH 7.4 (Ambion), 150 mM NaCl (Ambion), 1% Triton X-100 (Sigma-Aldrich), 0.1% sodium dodecyl sulfate (Promega) and 0.5% sodium deoxycholate (Sigma-Aldrich) with 1x HALT Protease and Phosphatase Inhibitor (Pierce). The DC Assay kit (Bio-Rad) was used to quantify protein concentration. The lysate was diluted in SDS Sample Buffer (125 mM Tris pH 6.8, 30% glycerol, 10% sodium dodecyl sulfate, 0.012% bromophenol blue) and denatured at 95 °C for 7 minutes. Between 20 and 40 micrograms of total protein was loaded per lane of 4-12% TGX Acrylamide Stain-Free gels (Bio-Rad). Prior to transfer by TransBlot Turbo (Bio-Rad), the Stain-Free gel was imaged to quantify total protein (Millipore or Bio-Rad). Blots were blocked in 5% milk or 5% bovine serum albumin in tris-buffered saline with tween prior to adding primary antibody: ADAR1 (Santa Cruz Biotechnology Cat# sc-73408, RRID:AB_2222767; Bethyl Cat# A303-883A, RRID:AB_2620233), ATF3 (Cell Signaling Technology Cat# 33593, RRID:AB_2799039), eIF2a (Abcam Cat# ab5369, RRID:AB_304838), eIF2a-Ser-51-P (Abcam Cat# ab32157, RRID:AB_732117), Phospho-NF-κB p65 (Ser 468) (Cell Signaling Technology Cat# 3039S, RRID:AB_330579), NF-κB p65 (Cell Signaling Technology Cat# 8242S, RRID:AB_10859369), beta-tubulin (Abcam Cat# ab6046, RRID:AB_2210370), cleaved PARP (Cell Signaling Technology Cat# 9541, RRID:AB_331426), GADD34 (Cell Signaling Technology Cat# 41222), Histone H3 (Abcam Cat# 10799, RRID:AB_470239), PACT (Cell Signaling Technology Cat# 13490, RRID AB_2798233), PKR (Cell Signaling Technology Cat# 3072, RRID:AB_2277600), PKR Thr-446-P (Abcam Cat# ab32036, RRID:AB_777310). Horseradish-peroxidase conjugated secondary antibodies (Jackson ImmunoResearch) and Clarity Western ECL Substrate (Bio-Rad) were used for detection via ChemiDoc (Bio-Rad). Image Lab (Bio-Rad) was used to determine band intensities which were normalized to total protein measured by imaging of the Stain-Free gel.

### Cell Viability and Crystal Violet Staining

For cell viability assessment, 5000 cells were plated in triplicate for each condition in opaque white 96-well plates. Cell viability was assessed by CellTiter-Glo 2.0 (Promega) per manufacturers protocol between three and four days after plating. For details on the number of cells plated and the timeline for cell viability assessment, see Supplemental Table 2.

For crystal violet staining, 2000 cells were plated per well of a 6-well dish. Between 15 to 20 days later, cells were washed briefly with 1x PBS prior to fixation in 100% methanol for 5 min. After drying, the cells were stained with 0.005% Crystal Violet solution containing 25% methanol (Sigma-Aldrich) prior to washing excess stain away with deionized water. Plates were imaged using a Bio-Rad ChemiDoc.

### Tumorigenesis

The Biological Evaluation Shared Resource at Purdue University Institute for Cancer research performed the tumorigenesis study. From the cranial end, the second left ventral mammary fat-pad of female NRG (NOD.Cg-Rag1tm1Mom Il2rgtm1Wjl/SzJ) mice, originally obtained from The Jackson Laboratory (RRID:IMSR_JAX:007799), were injected with 2.9 x 10^6^ HCC1806 cells suspended in equal volumes 1X PBS and Matrigel (Corning). Five days after injection, when tumors were palpable, mice were given acidified drinking water containing 2.4 mg/mL doxycycline. Tumor volume was measured manually using a caliper three days per week until the first mouse reached a humane euthanasia criterion. Five mice were injected per condition, one mouse injected with the sgNTA line was euthanized early due to poor health.

### RNA Purification and Analysis of rRNA Integrity

RNA was purified using the Nucleospin RNA kit (Macherey-Nagel). Ribosomal RNA integrity was determined using an Agilent TapeStation by the Genomics and Genome Editing Facility at Purdue University.

### RNA Sequencing and Analysis

Contaminating DNA was removed from total RNA by TURBO DNase (Thermo Fisher Scientific) prior to rRNA depletion using QIAseq® FastSelect™−rRNA HMR kit (Qiagen) per manufacturer’s protocol. RNA sequencing libraries were constructed with the NEBNext Ultra II RNA Library Preparation Kit for Illumina per manufacturer’s recommendations. Sequencing libraries were validated by Agilent Tapestation 4200 and quantified by Qubit 2.0 Fluorometer (ThermoFisher Scientific) as well as by quantitative PCR (KAPA Biosystems). The sequencing libraries were multiplexed and clustered onto a flowcell on the Illumina NovaSeq instrument. The samples were sequenced using a 2x150bp paired-end configuration. Image analysis and base calling were conducted by NovaSeq Control Software. Raw sequence data (.bcl files) generated from Illumina NovaSeq was converted into fastq files and de-multiplexed using Illumina bcl2fastq 2.20 software. One mismatch was allowed for index sequence identification.

Sequence reads were processed using Trimmomatic v.0.36 to remove adapter sequences and poor-quality nucleotides. Trimmed reads were aligned to GRCh38 reference genome using STAR aligner v2.5.2b (RRID:SCR_004463). Gene counts were determined using featureCounts from Subread v.1.5.2 (RRID:SCR_009803), only unique exonic reads were counted. Differential gene expression was determined using DESeq2 (RRID:SCR_015687, see Data Availability below for scripts) with shrunken fold changes using the ‘apeglm’ method ^49^. Gene set enrichment analysis was performed using ‘clusterProfiler’ (RRID:SCR_016884) with Gene Ongology (RRID:SCR_002811) terms from (Ashburner et al. 2000; Aleksander et al. 2023) or Hallmark gene sets from the Molecular Signatures Database (RRID:SCR_016863) (Liberzon et al. 2015). For Gene Set Enrichment Analysis with Hallmark gene sets, an additional gene set was included for previously identified ATF4 target genes (Wong et al. 2019).

### Quantitative PCR

LunaScript Supermix (NEB) was used to make cDNA for quantitative PCR (qPCR), using Luna Universal qPCR MasterMix (NEB) on a QuantStudio3 system (Thermo Scientific). All primers used for qPCR are listed in Supplemental Table 1. The amplification efficiency of each primer was verified to be within 90-110% allowing determination of ‘Fold Change’ by the ΔΔCt. Two reference genes were used for normalization, EEF1A1 and HSPA5, using their geometric mean Ct for calculating ΔCt.

### Analysis of TCGA data

For TCGA data, normalization of RNA-seq data, and z-scores calculations were performed as previously described ^35^. Breast cancer cell lines and TCGA tumor molecular subtypes were defined previously ^53^. The R packages RTCGA and survminer (RRID:SCR_021094) were used to determine breast cancer survival ^54,55^. The surv_cutpoint function of survminer was employed to determine an expression cutoff.

## Data Availability Statement

All analysis scripts are available at (https://github.com/cottrellka/Young_et_al_2025). Raw RNA-seq and gene count data is available at the Gene Expression Omnibus (GSE298233). Dependency (DepMap_Public_24Q4+Score,_Chronos, and Achilles+DRIVE+Marcotte,_DEMETER2), transcriptomic (Batch_corrected_Expression_Public_24Q4) and proteomic data for cancer cell lines (Harmonized_MS_CCLE_Gygi) were obtained from the DepMap portal (https://depmap.org/portal/download/custom/, RRID:SCR_017655)^56^. Transcriptomic data for TCGA BRCA samples (illuminahiseq_rnaseqv2-RSEM_genes) and clinical data (Merge_Clinical) were obtained from the Broad Institute FireBrowse and are available at http://firebrowse.org/.

## Acknowledgements

The data described here is in part based on data generated by the TCGA Research Network: https://www.cancer.gov/tcga. This work was supported by R00MD016946 (K.A. Cottrell), the Purdue Institute for Cancer Research (start-up funding, Robbers New Investigator Award to K.A. Cottrell, and support for the Biological Evaluation Shared Resource) NIH grant P30CA023168, and start-up funding from Purdue University Department of Biochemistry and College of Agriculture. Jill Hutchcroft, director of the Flow Cytometry and Cell Separation Facility at Purdue University for sorting iCas9 cell lines. Cottrell Lab rotation students Jolene Mach, Geethma Lirushie, Nima Goodarzi, Ayomide Adebesin, Yalan Huo, Troy Sievertsen, Tommy Sheeley and Reed Smith.

## Author Contributions

K.A.C. and A.A.Y. conceived the project. K.A.C., A.A.Y. and B.D.E. designed the experiments. A.A.Y., K.A.C., I.G.J., J.R.P., H.E.B., H.A.H., D.S.O., R.N.C., M.E.L., E.N.G., performed the experiments and/or provided materials. K.A.C, A.A.Y., I.G.J., J.R.P., H.E.B. performed the data analysis. K.A.C., A.A.Y., and I.G.J. wrote the manuscript. All authors edited the manuscript.

## References

Ahmad S, Zou T, Hwang J, Zhao L, Wang X, Davydenko A, Buchumenski I, Zhuang P, Fishbein AR, Capcha-Rodriguez D et al. 2025. PACT prevents aberrant activation of PKR by endogenous dsRNA without sequestration. Nat Commun 16: 3325.

Aleksander SA, Balhoff J, Carbon S, Cherry JM, Drabkin HJ, Ebert D, Feuermann M, Gaudet P, Harris NL, Hill DP et al. 2023. The Gene Ontology knowledgebase in 2023. Genetics 224.

Ashburner M, Ball CA, Blake JA, Botstein D, Butler H, Cherry JM, Davis AP, Dolinski K, Dwight SS, Eppig JT et al. 2000. Gene ontology: tool for the unification of biology. The Gene Ontology Consortium. Nat Genet 25: 25–29.

Bass BL. 2024. Adenosine deaminases that act on RNA, then and now. RNA 30: 521–529.

Bass BL, Weintraub H. 1988. An unwinding activity that covalently modifies its double-stranded RNA substrate. Cell **55**: 1089-1098.

Bianchini G, Balko JM, Mayer IA, Sanders ME, Gianni L. 2016. Triple-negative breast cancer: challenges and opportunities of a heterogeneous disease. Nat Rev Clin Oncol 13: 674–690.

Bonnet MC, Daurat C, Ottone C, Meurs EF. 2006. The N-terminus of PKR is responsible for the activation of the NF-kappaB signaling pathway by interacting with the IKK complex. Cell Signal 18: 1865–1875.

Bonnet MC, Weil R, Dam E, Hovanessian AG, Meurs EF. 2000. PKR stimulates NF-kappaB irrespective of its kinase function by interacting with the IkappaB kinase complex. Mol Cell Biol 20: 4532–4542.

Broad D. 2024a. DepMap 24Q2 Public.

Broad D. 2024b. DepMap 24Q4 Public.

Cao J, Wu L, Zhang SM, Lu M, Cheung WK, Cai W, Gale M, Xu Q, Yan Q. 2016. An easy and efficient inducible CRISPR/Cas9 platform with improved specificity for multiple gene targeting. Nucleic Acids Res 44: e149.

Chakrabarti A, Jha BK, Silverman RH. 2011. New insights into the role of RNase L in innate immunity. J Interferon Cytokine Res 31: 49–57.

Chen DS, Mellman I. 2017. Elements of cancer immunity and the cancer-immune set point. Nature 541: 321–330.

Chen R, Ishak CA, De Carvalho DD. 2021. Endogenous Retroelements and the Viral Mimicry Response in Cancer Therapy and Cellular Homeostasis. Cancer Discov 11: 2707–2725.

Chen YG, Hur S. 2022. Cellular origins of dsRNA, their recognition and consequences. Nat Rev Mol Cell Biol 23: 286–301.

Chukwurah E, Farabaugh KT, Guan BJ, Ramakrishnan P, Hatzoglou M. 2021. A tale of two proteins: PACT and PKR and their roles in inflammation. FEBS J 288: 6365–6391.

Chung H, Calis JJA, Wu X, Sun T, Yu Y, Sarbanes SL, Dao Thi VL, Shilvock AR, Hoffmann HH, Rosenberg BR et al. 2018. Human ADAR1 Prevents Endogenous RNA from Triggering Translational Shutdown. Cell 172: 811–824.e814.

Clerzius G, Shaw E, Daher A, Burugu S, Gélinas JF, Ear T, Sinck L, Routy JP, Mouland AJ, Patel RC et al. 2013. The PKR activator, PACT, becomes a PKR inhibitor during HIV-1 replication. Retrovirology **10**: 96.

Costa-Mattioli M, Walter P. 2020. The integrated stress response: From mechanism to disease. Science 368.

Cottrell KA, Andrews RJ, Bass BL. 2024a. The competitive landscape of the dsRNA world. Mol Cell 84: 107–119.

Cottrell KA, Ryu S, Pierce JR, Soto Torres L, Bohlin HE, Schab AM, Weber JD. 2024b. Induction of Viral Mimicry Upon Loss of DHX9 and ADAR1 in Breast Cancer Cells. Cancer Res Commun 4: 986–1003.

Curigliano G, Goldhirsch A. 2011. The triple-negative subtype: new ideas for the poorest prognosis breast cancer. J Natl Cancer Inst Monogr 2011: 108–110.

Dempster JM, Rossen J, Kazachkova M, Pan J, Kugener G, Root DE, Tsherniak A. 2019. Extracting Biological Insights from the Project Achilles Genome-Scale CRISPR Screens in Cancer Cell Lines. bioRxiv: 720243.

Dickerman BK, White CL, Kessler PM, Sadler AJ, Williams BR, Sen GC. 2015. The protein activator of protein kinase R, PACT/RAX, negatively regulates protein kinase R during mouse anterior pituitary development. FEBS J 282: 4766–4781.

Doench JG, Fusi N, Sullender M, Hegde M, Vaimberg EW, Donovan KF, Smith I, Tothova Z, Wilen C, Orchard R et al. 2016. Optimized sgRNA design to maximize activity and minimize off-target effects of CRISPR-Cas9. Nat Biotechnol 34: 184–191.

Elbarbary RA, Li W, Tian B, Maquat LE. 2013. STAU1 binding 3’ UTR IRAlus complements nuclear retention to protect cells from PKR-mediated translational shutdown. Genes Dev 27: 1495–1510.

Farabaugh KT, Krokowski D, Guan BJ, Gao Z, Gao XH, Wu J, Jobava R, Ray G, de Jesus TJ, Bianchi MG et al. 2020. PACT-mediated PKR activation acts as a hyperosmotic stress intensity sensor weakening osmoadaptation and enhancing inflammation. Elife 9.

Gal-Ben-Ari S, Barrera I, Ehrlich M, Rosenblum K. 2018. PKR: A Kinase to Remember. Front Mol Neurosci 11: 480.

Gannon HS, Zou T, Kiessling MK, Gao GF, Cai D, Choi PS, Ivan AP, Buchumenski I, Berger AC, Goldstein JT et al. 2018. Identification of ADAR1 adenosine deaminase dependency in a subset of cancer cells. Nat Commun 9: 5450.

Gil J, Alcamí J, Esteban M. 1999. Induction of apoptosis by double-stranded-RNA-dependent protein kinase (PKR) involves the alpha subunit of eukaryotic translation initiation factor 2 and NF-kappaB. Mol Cell Biol 19: 4653–4663.

Gil J, Rullas J, García MA, Alcamí J, Esteban M. 2001. The catalytic activity of dsRNA-dependent protein kinase, PKR, is required for NF-kappaB activation. Oncogene 20: 385–394.

Guirguis AA, Ofir-Rosenfeld Y, Knezevic K, Blackaby W, Hardick D, Chan YC, Motazedian A, Gillespie A, Vassiliadis D, Lam EYN et al. 2023. Inhibition of METTL3 Results in a Cell-Intrinsic Interferon Response That Enhances Antitumor Immunity. Cancer Discov 13: 2228–2247.

Heyam A, Coupland CE, Dégut C, Haley RA, Baxter NJ, Jakob L, Aguiar PM, Meister G, Williamson MP, Lagos D et al. 2017. Conserved asymmetry underpins homodimerization of Dicer-associated double-stranded RNA-binding proteins. Nucleic Acids Res 45: 12577–12584.

Hovanessian AG, Justesen J. 2007. The human 2’-5’oligoadenylate synthetase family: unique interferon-inducible enzymes catalyzing 2’-5’ instead of 3’-5’ phosphodiester bond formation. Biochimie 89: 779–788.

Hu SB, Heraud-Farlow J, Sun T, Liang Z, Goradia A, Taylor S, Walkley CR, Li JB. 2023. ADAR1p150 prevents MDA5 and PKR activation via distinct mechanisms to avert fatal autoinflammation. Mol Cell 83: 3869–3884.e3867.

Huang HT, Seo HS, Zhang T, Wang Y, Jiang B, Li Q, Buckley DL, Nabet B, Roberts JM, Paulk J et al. 2017. MELK is not necessary for the proliferation of basal-like breast cancer cells. Elife 6.

Huang W, Zhu Q, Shi Z, Tu Y, Li Q, Zheng W, Yuan Z, Li L, Zu X, Hao Y et al. 2024. Dual inhibitors of DNMT and HDAC induce viral mimicry to induce antitumour immunity in breast cancer. Cell Death Discov 10: 143.

Ishizuka JJ, Manguso RT, Cheruiyot CK, Bi K, Panda A, Iracheta-Vellve A, Miller BC, Du PP, Yates KB, Dubrot J et al. 2019. Loss of ADAR1 in tumours overcomes resistance to immune checkpoint blockade. Nature 565: 43–48.

Jeffrey IW, Kadereit S, Meurs EF, Metzger T, Bachmann M, Schwemmle M, Hovanessian AG, Clemens MJ. 1995. Nuclear localization of the interferon-inducible protein kinase PKR in human cells and transfected mouse cells. Exp Cell Res 218: 17–27.

Jiang X, Muthusamy V, Fedorova O, Kong Y, Kim DJ, Bosenberg M, Pyle AM, Iwasaki A. 2019. Intratumoral delivery of RIG-I agonist SLR14 induces robust antitumor responses. J Exp Med 216: 2854–2868.

Kim Y, Lee JH, Park JE, Cho J, Yi H, Kim VN. 2014. PKR is activated by cellular dsRNAs during mitosis and acts as a mitotic regulator. Genes Dev 28: 1310–1322.

Kumar A, Haque J, Lacoste J, Hiscott J, Williams BR. 1994. Double-stranded RNA-dependent protein kinase activates transcription factor NF-kappa B by phosphorylating I kappa B. Proc Natl Acad Sci U S A 91: 6288–6292.

Kung CP, Cottrell KA, Ryu S, Bramel ER, Kladney RD, Bao EA, Freeman EC, Sabloak T, Maggi L, Weber JD. 2021. Evaluating the therapeutic potential of ADAR1 inhibition for triple-negative breast cancer. Oncogene 40: 189–202.

Li S, Peters GA, Ding K, Zhang X, Qin J, Sen GC. 2006. Molecular basis for PKR activation by PACT or dsRNA. Proc Natl Acad Sci U S A 103: 10005–10010.

Liberzon A, Birger C, Thorvaldsdóttir H, Ghandi M, Mesirov JP, Tamayo P. 2015. The Molecular Signatures Database (MSigDB) hallmark gene set collection. Cell Syst 1: 417–425.

Liddicoat BJ, Piskol R, Chalk AM, Ramaswami G, Higuchi M, Hartner JC, Li JB, Seeburg PH, Walkley CR. 2015. RNA editing by ADAR1 prevents MDA5 sensing of endogenous dsRNA as nonself. Science 349: 1115–1120.

Liu H, Golji J, Brodeur LK, Chung FS, Chen JT, deBeaumont RS, Bullock CP, Jones MD, Kerr G, Li L, et al. 2019. Tumor-derived IFN triggers chronic pathway agonism and sensitivity to ADAR loss. Nat Med 25: 95–102.

Lu T, Ma P, Fang H, Chen A, Xu J, Kuang X, Wang M, Su L, Wang S, Zhang Y et al. 2025. Prkra dimer senses double-stranded RNAs to dictate global translation efficiency. Mol Cell 85: 2032–2047.e2039.

Manjunath L, Santiago G, Ortega P, Sanchez A, Oh S, Garcia A, Li J, Duong D, Bournique E, Bouin A et al. 2025. Cooperative role of PACT and ADAR1 in preventing aberrant PKR activation by self-derived double-stranded RNA. Nat Commun 16: 3246.

Matsumoto M, Oshiumi H, Seya T. 2011. Antiviral responses induced by the TLR3 pathway. Rev Med Virol 21: 67–77.

McFarland JM, Ho ZV, Kugener G, Dempster JM, Montgomery PG, Bryan JG, Krill-Burger JM, Green TM, Vazquez F, Boehm JS et al. 2018. Improved estimation of cancer dependencies from large-scale RNAi screens using model-based normalization and data integration. Nat Commun 9: 4610.

Mendoza HG, Beal PA. 2024. Structural and functional effects of inosine modification in mRNA. RNA 30: 512–520.

Meyer C, Garzia A, Mazzola M, Gerstberger S, Molina H, Tuschl T. 2018. The TIA1 RNA-Binding Protein Family Regulates EIF2AK2-Mediated Stress Response and Cell Cycle Progression. Mol Cell 69: 622–635.e626.

Morad G, Helmink BA, Sharma P, Wargo JA. 2021. Hallmarks of response, resistance, and toxicity to immune checkpoint blockade. Cell 184: 5309–5337.

Novoa I, Zeng H, Harding HP, Ron D. 2001. Feedback inhibition of the unfolded protein response by GADD34-mediated dephosphorylation of eIF2alpha. J Cell Biol 153: 1011–1022.

Park H, Davies MV, Langland JO, Chang HW, Nam YS, Tartaglia J, Paoletti E, Jacobs BL, Kaufman RJ, Venkatesan S. 1994. TAR RNA-binding protein is an inhibitor of the interferon-induced protein kinase PKR. Proc Natl Acad Sci U S A 91: 4713–4717.

Patel CV, Handy I, Goldsmith T, Patel RC. 2000. PACT, a stress-modulated cellular activator of interferon-induced double-stranded RNA-activated protein kinase, PKR. J Biol Chem 275: 37993–37998.

Patel RC, Sen GC. 1998. PACT, a protein activator of the interferon-induced protein kinase, PKR. EMBO J 17: 4379–4390.

Pestal K, Funk CC, Snyder JM, Price ND, Treuting PM, Stetson DB. 2015. Isoforms of RNA-Editing Enzyme ADAR1 Independently Control Nucleic Acid Sensor MDA5-Driven Autoimmunity and Multi-organ Development. Immunity 43: 933–944.

Peters GA, Dickerman B, Sen GC. 2009. Biochemical analysis of PKR activation by PACT. Biochemistry 48: 7441–7447.

Peters GA, Hartmann R, Qin J, Sen GC. 2001. Modular structure of PACT: distinct domains for binding and activating PKR. Mol Cell Biol 21: 1908–1920.

Rehwinkel J, Gack MU. 2020. RIG-I-like receptors: their regulation and roles in RNA sensing. Nat Rev Immunol 20: 537–551.

Singh M, Castillo D, Patel CV, Patel RC. 2011. Stress-induced phosphorylation of PACT reduces its interaction with TRBP and leads to PKR activation. Biochemistry 50: 4550–4560.

Sinigaglia K, Cherian A, Du Q, Lacovich V, Vukić D, Melicherová J, Linhartova P, Zerad L, Stejskal S, Malik R et al. 2024. An ADAR1 dsRBD3-PKR kinase domain interaction on dsRNA inhibits PKR activation. Cell Rep 43: 114618.

Stewart SA, Dykxhoorn DM, Palliser D, Mizuno H, Yu EY, An DS, Sabatini DM, Chen IS, Hahn WC, Sharp PA et al. 2003. Lentivirus-delivered stable gene silencing by RNAi in primary cells. RNA 9: 493–501.

Stringer BW, Day BW, D’Souza RCJ, Jamieson PR, Ensbey KS, Bruce ZC, Lim YC, Goasdoué K, Offenhäuser C, Akgül S et al. 2019. A reference collection of patient-derived cell line and xenograft models of proneural, classical and mesenchymal glioblastoma. Sci Rep 9: 4902.

Takahashi T, Miyakawa T, Zenno S, Nishi K, Tanokura M, Ui-Tei K. 2013. Distinguishable in vitro binding mode of monomeric TRBP and dimeric PACT with siRNA. PLoS One 8: e63434.

Valente L, Nishikura K. 2007. RNA binding-independent dimerization of adenosine deaminases acting on RNA and dominant negative effects of nonfunctional subunits on dimer functions. J Biol Chem 282: 16054–16061.

Waks AG, Winer EP. 2019. Breast Cancer Treatment: A Review. JAMA 321: 288–300.

Wong YL, LeBon L, Basso AM, Kohlhaas KL, Nikkel AL, Robb HM, Donnelly-Roberts DL, Prakash J, Swensen AM, Rubinstein ND et al. 2019. eIF2B activator prevents neurological defects caused by a chronic integrated stress response. Elife 8.

Young AA, Bohlin HE, Pierce JR, Cottrell KA. 2024. Suppression of double-stranded RNA sensing in cancer: molecular mechanisms and therapeutic potential. Biochem Soc Trans.

Zamanian-Daryoush M, Mogensen TH, DiDonato JA, Williams BR. 2000. NF-kappaB activation by double-stranded-RNA-activated protein kinase (PKR) is mediated through NF-kappaB-inducing kinase and IkappaB kinase. Mol Cell Biol 20: 1278–1290.

Zou T, Zhou M, Gupta A, Zhuang P, Fishbein AR, Wei HY, Capcha-Rodriguez D, Zhang Z, Cherniack AD, Meyerson M. 2024. XRN1 deletion induces PKR-dependent cell lethality in interferon-activated cancer cells. Cell Rep 43: 113600.

