## Supplemental Information for "PACT suppresses PKR activation through dsRNA binding and dimerization, and is a therapeutic target for triple-negative breast cancer"

**Contents:**

Supplemental Methods
Coding and amino acid sequences of wild-type, mutant and truncated PACT constructs
Figures S1-S5
Uncropped blots for main and supplemental figures

#### Supplemental Methods

##### Immunofluorescence

HCC1806 cells with expression of the indicated constructs were plated on glass coverslips (Corning) in 6-well dishes. After two days, the cells were washed in 1X PBS prior to fixation with 4% paraformaldehyde (Thermo-Scientific) in 1X PBS for 15 minutes at room temperature. Subsequently, the coverslips were washed in 1X PBS and the cells permeabilized with 0.15% Triton X-100 (Fisher) in 1X PBS. The coverslips were blocked in Protein Block (Aligent/Dako, X090930-2) prior to binding of primary antibodies for PKR (Cell Signaling Technology Cat# 3072, RRID:AB_2277600) and PACT (Santa Cruz Biotechnology Cat# sc-377103, RRID:AB_ 3665861), washing in 1X PBS and binding of secondary antibodies (Thermo Scientific, A21207 (RRID:AB_141637), A21203 (RRID:AB_2535789)). All antibodies were diluted in Antibody Diluent (Agilent/Dako, S302283-2). DAPI (Sigma-Aldrich) was added to the diluted secondary antibody during binding. After secondary antibody binding, the coverslips were washed in PBS and once in water before mounting on glass slides with Vectashield Antifade Mounting Media (Vector Laboratories). Fluorescence microscopy images were obtained with a DM6 B microscope (Leica) with a 20x objective and a DFC90000 GT monochrome digital camera (Leica). Fluorescence images were captured with Leica Application Suite X (Leica, RRID:SCR_013673) and resized and formatted with Fiji (RRID:SCR_002285).

**Coding sequences for PACT constructs**

>PACT_wobble_CDS
ATGTCCCAGAGCAGGCACCGCGCCGAGGCCCCGCCGCTGGAGCGCGAGGACAGTGGGACCTTCAGTTT
GGGGAAGATGATAACAGCTAAGCCAGGGAAAACACCGATTCAGGTATTACACGAATACGGCATGAAGACC
AAGAACATCCCAGTTTATGAATGTGAAAGATCTGATGTGCAAATACACGTGCCCACTTTCACCTTCAGAGT
AACCGTgGGcGAtATcACaTGtAC*cGG*TGAAGGTACAAGTAAGAAGCTcGCcAAGCAccGcGCaGCcG*AaG*CT
GCCATAAACATTTTGAAAGCCAATGCAAGTATTTGCTTTGCAGTTCCTGACCCCTTAATGCCTGACCCTTC
CAAGCAACCAAAGAACCAGCTTAATCCTATTGGTTCATTACAGGAATTGGCTATTCATCATGGCTGGAGAC
TTCCTGAATATACCCTTTCCCAGGAGGGAGGACCTGCTCATAAGAGAGAATATACTACAATTTGCAGGCTA
GAGTCATTTATGGAAACTGGAAAGGGGGCATCAAAAAAGCAAGCCAAAAGGAATGCTGCTGAGAAATTTC
TTGCCAAATTTAGTAATATTTCTCCAGAGAACCACATTTCTTTAACAAATGTAGTAGGACATTCTTTAGGAT
GTACTTGGCATTCCTTGAGGAATTCTCCTGGTGAAAAGATCAACTTACTGAAAAGAAGCCTCCTTAGTATT
CCAAATACAGATTACATCCAGCTGCTTAGTGAAATTGCCAAGGAACAAGGTTTTAATATAACATATTTGGAT
ATAGATGAACTGAGCGCCAATGGACAATATCAATGTCTTGCTGAACTGTCCACCAGCCCCATCACAGTCT
GTCATGGCTCCGGTATCTCCTGTGGCAATGCACAAAGTGATGCAGCTCACAATGCTTTGCAGTATTTAAAG
ATAATAGCAGAAAGAAAGTAA

-underlined bases are the target sites of sgPACT-1 and sgPACT-2, lowercase bases are wobble mutations to prevent base-pairing with the sgRNA or to disrupt the PAM (italicized)

>PACT_wobble_AA-mutant_CDS
ATGTCCCAGAGCAGGCACCGCGCCGAGGCCCCGCCGCTGGAGCGCGAGGACAGTGGGACCTTCAGTTT
GGGGAAGATGATAACAGCTAAGCCAGGGAAAACACCGATTCAGGTATTACACGAATACGGCATGAAGACC
AAGAACATCCCAGTTTATGAATGTGAAAGATCTGATGTGCAAATACACGTGCCCACTTTCACCTTCAGAGT
AACCGTGGGCGATATCACATGTACCGGTGAAGGTACAAGT**gcggca**CTCGCCAAGCACCGCGCAGCCGAA
GCTGCCATAAACATTTTGAAAGCCAATGCAAGTATTTGCTTTGCAGTTCCTGACCCCTTAATGCCTGACCC
TTCCAAGCAACCAAAGAACCAGCTTAATCCTATTGGTTCATTACAGGAATTGGCTATTCATCATGGCTGGA
GACTTCCTGAATATACCCTTTCCCAGGAGGGAGGACCTGCTCATAAGAGAGAATATACTACAATTTGCAGG
CTAGAGTCATTTATGGAAACTGGAAAGGGGGCATCA**gcagcg**CAAGCCaaaAGGAATGCTGCTGAGAAATTT
CTTGCCAAATTTAGTAATATTTCTCCAGAGAACCACATTTCTTTAACAAATGTAGTAGGACATTCTTTAGGAT
GTACTTGGCATTCCTTGAGGAATTCTCCTGGTGAAAAGATCAACTTACTGAAAAGAAGCCTCCTTAGTATT
CCAAATACAGATTACATCCAGCTGCTTAGTGAAATTGCCAAGGAACAAGGTTTTAATATAACATATTTGGAT
ATAGATGAACTGAGCGCCAATGGACAATATCAATGTCTTGCTGAACTGTCCACCAGCCCCATCACAGTCT
GTCATGGCTCCGGTATCTCCTGTGGCAATGCACAAAGTGATGCAGCTCACAATGCTTTGCAGTATTTAAAG
ATAATAGCAGAAAGAAAGTAA

-lowercase boldened bases are mutations to disrupt dsRNA binding

>PACT_wobble_EAA-dsRBD-mutant_CDS
ATGTCCCAGAGCAGGCACCGCGCCGAGGCCCCGCCGCTGGAGCGCGAGGACAGTGGGACCTTCAGTTT
GGGGAAGATGATAACAGCTAAGCCAGGGAAAACACCGATTCAGGTATTACACGAATACGGCATGAAGACC
AAGAACATCCCAGTTTATGAATGTGAAAGATCTGATGTGCAAATACACGTGCCCACTTTCACCTTCAGAGT
AACCGTGGGCGATATCACATGTACCGGTGAAGGTACAAGT**gaagcg**CTCGCC**gcg**CACCGCGCAGCCGAA
GCTGCCATAAACATTTTGAAAGCCAATGCAAGTATTTGCTTTGCAGTTCCTGACCCCTTAATGCCTGACCC
TTCCAAGCAACCAAAGAACCAGCTTAATCCTATTGGTTCATTACAGGAATTGGCTATTCATCATGGCTGGA
GACTTCCTGAATATACCCTTTCCCAGGAGGGAGGACCTGCTCATAAGAGAGAATATACTACAATTTGCAGG
CTAGAGTCATTTATGGAAACTGGAAAGGGGGCATCA**gaagcg**CAAGCC**gca**AGGAATGCTGCTGAGAAATTT
CTTGCCAAATTTAGTAATATTTCTCCAGAGAACCACATTTCTTTAACAAATGTAGTAGGACATTCTTTAGGAT

GTACTTGGCATTCCTTGAGGAATTCTCCTGGTGAAAAGATCAACTTACTGAAAAGAAGCCTCCTTAGTATT
CCAAATACAGATTACATCCAGCTGCTTAGTGAAATTGCCAAGGAACAAGGTTTTAATATAACATATTTGGAT
ATAGATGAACTGAGCGCCAATGGACAATATCAATGTCTTGCTGAACTGTCCACCAGCCCCATCACAGTCT
GTCATGGCTCCGGTATCTCCTGTGGCAATGCACAAAGTGATGCAGCTCACAATGCTTTGCAGTATTTAAAG
ATAATAGCAGAAAGAAAGTAA

-lowercase boldened bases are mutations to disrupt dsRNA binding

>PACT_wobble_Δd3_CDS
ATGTCCCAGAGCAGGCACCGCGCCGAGGCCCCGCCGCTGGAGCGCGAGGACAGTGGGACCTTCAGTTT
GGGGAAGATGATAACAGCTAAGCCAGGGAAAACACCGATTCAGGTATTACACGAATACGGCATGAAGACC
AAGAACATCCCAGTTTATGAATGTGAAAGATCTGATGTGCAAATACACGTGCCCACTTTCACCTTCAGAGT
AACCGTGGGCGATATCACATGTACCGGTGAAGGTACAAGTAAGAAGCTCGCCAAGCACCGCGCAGCCGA
AGCTGCCATAAACATTTTGAAAGCCAATGCAAGTATTTGCTTTGCAGTTCCTGACCCCTTAATGCCTGACC
CTTCCAAGCAACCAAAGAACCAGCTTAATCCTATTGGTTCATTACAGGAATTGGCTATTCATCATGGCTGG
AGACTTCCTGAATATACCCTTTCCCAGGAGGGAGGACCTGCTCATAAGAGAGAATATACTACAATTTGCAG
GCTAGAGTCATTTATGGAAACTGGAAAGGGGGCATCAAAAAAGCAAGCCAAAAGGAATGCTGCTGAGAAA
TTTCTTGCCAAATTTAGTAATATTTCTCCAGAGAACCACATTTCTTTAACAAATGTAGTAGGACATTCTTTAG
GATGTACTTGGCATTCCTTGAGgaattcGTAA

-lowercase bases are the EcoRI site used to truncate PACT

>PACT_wobble_Δd3-GST_CDS
ATGTCCCAGAGCAGGCACCGCGCCGAGGCCCCGCCGCTGGAGCGCGAGGACAGTGGGACCTTCAGTTT
GGGGAAGATGATAACAGCTAAGCCAGGGAAAACACCGATTCAGGTATTACACGAATACGGCATGAAGACC
AAGAACATCCCAGTTTATGAATGTGAAAGATCTGATGTGCAAATACACGTGCCCACTTTCACCTTCAGAGT
AACCGTGGGCGATATCACATGTACCGGTGAAGGTACAAGTAAGAAGCTCGCCAAGCACCGCGCAGCCGA
AGCTGCCATAAACATTTTGAAAGCCAATGCAAGTATTTGCTTTGCAGTTCCTGACCCCTTAATGCCTGACC
CTTCCAAGCAACCAAAGAACCAGCTTAATCCTATTGGTTCATTACAGGAATTGGCTATTCATCATGGCTGG
AGACTTCCTGAATATACCCTTTCCCAGGAGGGAGGACCTGCTCATAAGAGAGAATATACTACAATTTGCAG
GCTAGAGTCATTTATGGAAACTGGAAAGGGGGCATCAAAAAAGCAAGCCAAAAGGAATGCTGCTGAGAAA
TTTCTTGCCAAATTTAGTAATATTTCTCCAGAGAACCACATTTCTTTAACAAATGTAGTAGGACATTCTTTAG
GATGTACTTGGCATTCCTTGAGgaattcACATATGTCCCCTATACTAGGTTATTGGAAAATTAAGGGCCTTGT
GCAACCCACTCGACTTCTTTTGGAATATCTTGAAGAAAAATATGAAGAGCATTTGTATGAGCGCGATGAAG
GTGATAAATGGCGAAACAAAAAGTTTGAATTGGGTTTGGAGTTTCCCAATCTTCCTTATTATATTGATGGTG
ATGTTAAATTAACACAGTCTATGGCCATCATACGTTATATAGCTGACAAGCACAACATGTTGGGTGGTTGT
CCAAAAGAGCGTGCAGAGATTTCAATGCTTGAAGGAGCGGTTTTGGATATTAGATACGGTGTTTCGAGAAT
TGCATATAGTAAAGACTTTGAAACTCTCAAAGTTGATTTTCTTAGCAAGCTACCTGAAATGCTGAAAATGTT
CGAAGATCGTTTATGTCATAAAACATATTTAAATGGTGATCATGTAACCCATCCTGACTTCATGTTGTATGA
CGCTCTTGATGTTGTTTTATACATGGACCCAATGTGCCTGGATGCGTTCCCAAAATTAGTTTGTTTTAAAAA
ACGTATTGAAGCTATCCCACAAATTGATAAGTACTTGAAATCCAGCAAGTATATAGCATGGCCTTTGCAGG
GCTGGCAAGCCACGTTTGGTGGTGGCGACCATCCTCCAAAATGA

-lowercase bases are the EcoRI site used to truncate PACT and add GST

**Amino acid sequences of PACT constructs**

>PACT_WT

MSQSRHRAEAPPLEREDSGTFSLGKMITAKPGKTPIQVLHEYGMKTKNIPVYECERSDVQIHVPTFTFRVTVGD
ITCTGEGTSKKLAKHRAAEAAINILKANASICFAVPDPLMPDPSKQPKNQLNPIGSLQELAIHHGWRLPEYTLSQ
EGGPAHKREYTTICRLESFMETGKGASKKQAKRNAAEKFLAKFSNISPENHISLTNVVGHSLGCTWHSLRNSP
GEKINLLKRSLLSIPNTDYIQLLSEIAKEQGFNITYLDIDELSANGQYQCLAELSTSPITVCHGSGISCGNAQSDAA
HNALQYLKIIAERK

>PACT_AA-dsRBD-mutant
MSQSRHRAEAPPLEREDSGTFSLGKMITAKPGKTPIQVLHEYGMKTKNIPVYECERSDVQIHVPTFTFRVTVGD
ITCTGEGTSAALAKHRAAEAAINILKANASICFAVPDPLMPDPSKQPKNQLNPIGSLQELAIHHGWRLPEYTLSQ
EGGPAHKREYTTICRLESFMETGKGASAAQAKRNAAEKFLAKFSNISPENHISLTNVVGHSLGCTWHSLRNSP
GEKINLLKRSLLSIPNTDYIQLLSEIAKEQGFNITYLDIDELSANGQYQCLAELSTSPITVCHGSGISCGNAQSDAA
HNALQYLKIIAERK

>PACT-EAA-dsRBD-mutant
MSQSRHRAEAPPLEREDSGTFSLGKMITAKPGKTPIQVLHEYGMKTKNIPVYECERSDVQIHVPTFTFRVTVGD
ITCTGEGTSEALAAHRAAEAAINILKANASICFAVPDPLMPDPSKQPKNQLNPIGSLQELAIHHGWRLPEYTLSQ
EGGPAHKREYTTICRLESFMETGKGASEAQAARNAAEKFLAKFSNISPENHISLTNVVGHSLGCTWHSLRNSP
GEKINLLKRSLLSIPNTDYIQLLSEIAKEQGFNITYLDIDELSANGQYQCLAELSTSPITVCHGSGISCGNAQSDAA
HNALQYLKIIAERK

>PACT-Δd3
MSQSRHRAEAPPLEREDSGTFSLGKMITAKPGKTPIQVLHEYGMKTKNIPVYECERSDVQIHVPTFTFRVTVGD
ITCTGEGTSKKLAKHRAAEAAINILKANASICFAVPDPLMPDPSKQPKNQLNPIGSLQELAIHHGWRLPEYTLSQ
EGGPAHKREYTTICRLESFMETGKGASKKQAKRNAAEKFLAKFSNISPENHISLTNVVGHSLGCTWHSLRNS

>PACT-Δd3-GST
MSQSRHRAEAPPLEREDSGTFSLGKMITAKPGKTPIQVLHEYGMKTKNIPVYECERSDVQIHVPTFTFRVTVGD
ITCTGEGTSKKLAKHRAAEAAINILKANASICFAVPDPLMPDPSKQPKNQLNPIGSLQELAIHHGWRLPEYTLSQ
EGGPAHKREYTTICRLESFMETGKGASKKQAKRNAAEKFLAKFSNISPENHISLTNVVGHSLGCTWHSLRNSH
MSPILGYWKIKGLVQPTRLLLEYLEEKYEEHLYERDEGDKWRNKKFELGLEFPNLPYYIDGDVKLTQSMAIIRYI
ADKHNMLGGCPKERAEISMLEGAVLDIRYGVSRIAYSKDFETLKVDFLSKLPEMLKMFEDRLCHKTYLNGDHVT
HPDFMLYDALDVVLYMDPMCLDAFPKLVCFKKRIEAIPQIDKYLKSSKYIAWPLQGWQATFGGGDHPPK

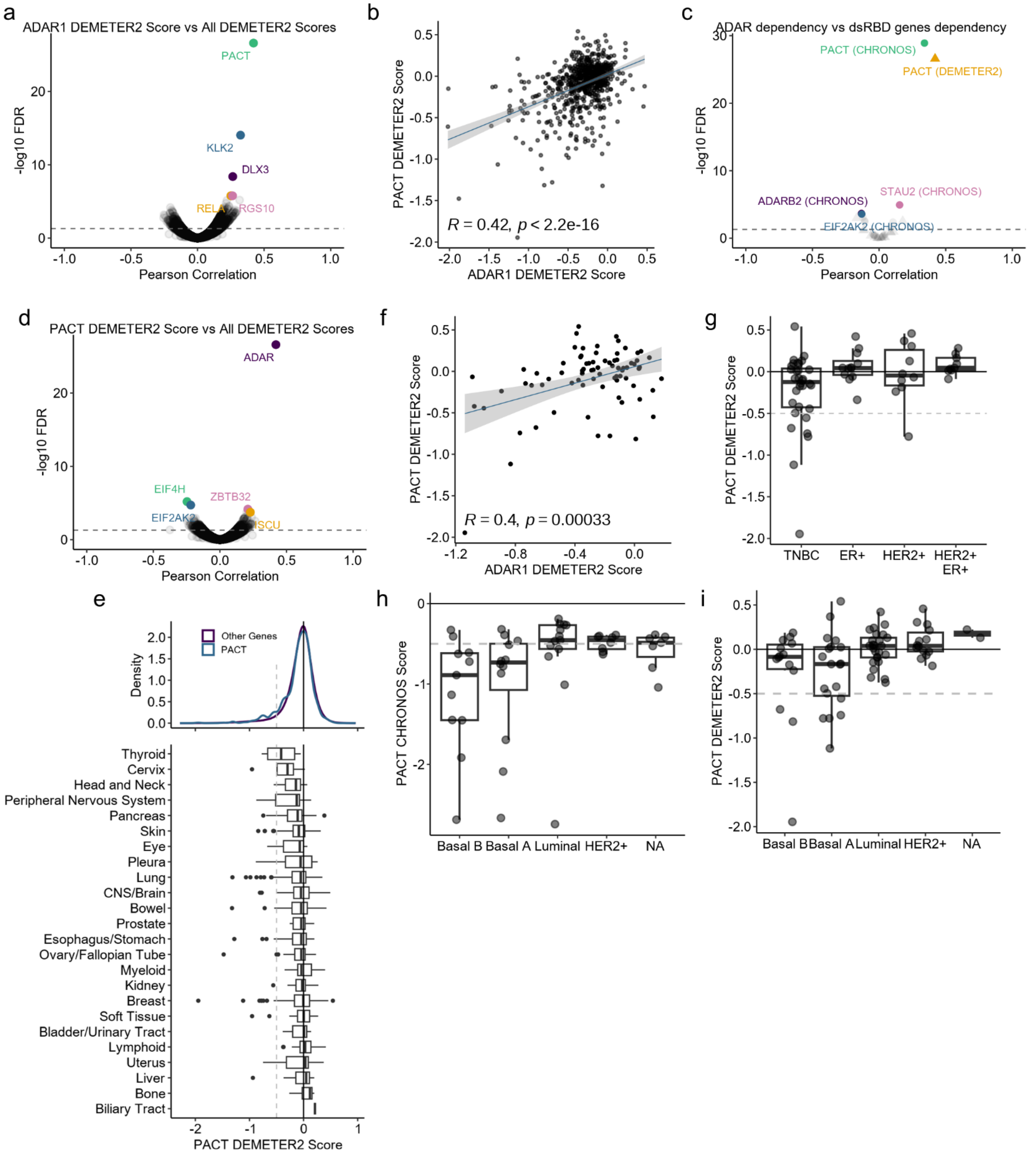


**Figure S1**

**a** and **d** Volcano plots of Pearson correlation coefficients and FDR corrected p-values for pairwise comparisons between ADAR1 DEMETER2 score (**a**) or PACT DEMETER2 score (**d**) and DEMETER2 scores for all genes in DepMap across all cell lines. **b** Correlation between PACT and ADAR1 DEMETER2 scores for all DepMap cell lines, Pearson correlation coefficient and p-value shown. **c** Volcano plot of Pearson correlation coefficients and FDR corrected p-values for pairwise comparisons between ADAR1 DEMETER2 or CHRONOS score and DEMETER2 or CHRONOS scores for all dsRBD containing genes in DepMap across all cell lines. **e** Top, density plot of DEMTER2 scores for PACT or all other genes. Bottom, boxplots for PACT DEMTER2 score by lineage. **f** Correlation between PACT and ADAR1 DEMETER2 scores for breast cancer cell lines, Pearson correlation coefficient and p-value shown. **g-i** Boxplot of PACT CHRONOS or DEMETER2 scores of breast cancer cell lines separated by subtype. All data shown from DepMap.

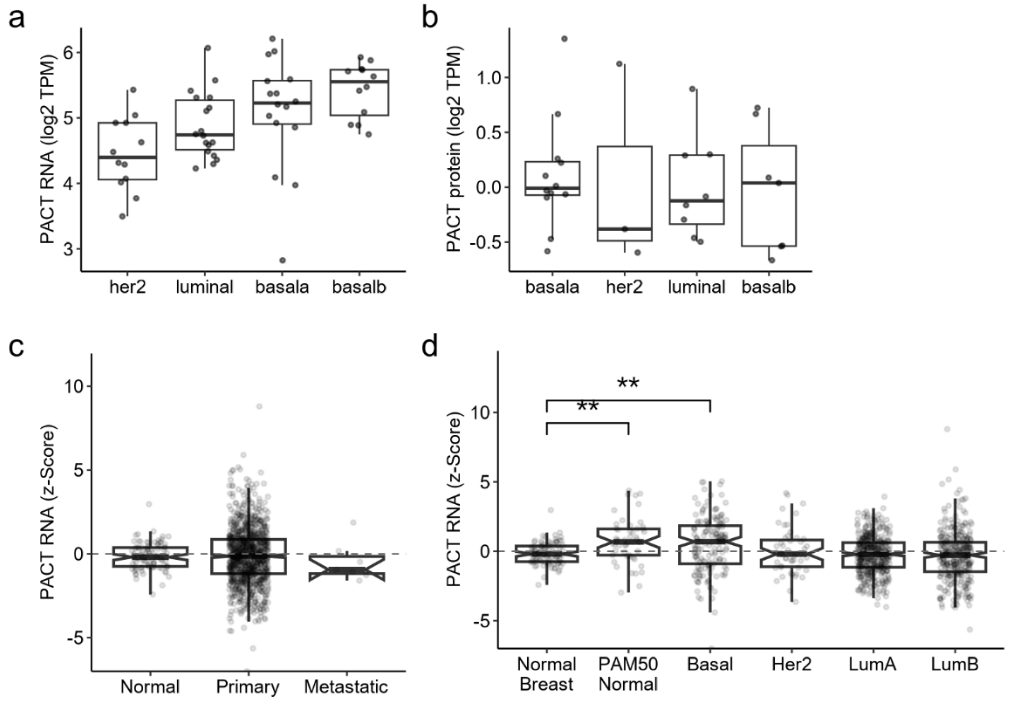


**Figure S2**

Expression of PACT at the RNA (**a**) or protein level (**b**) in breast cancer cell lines separated by subtype. **c** - **d** Expression of PACT at the RNA level in normal and breast tumor samples separated by subtypes or tumor location. For panels **a**-**b**, data from DepMap; **c**-**d** from TCGA.

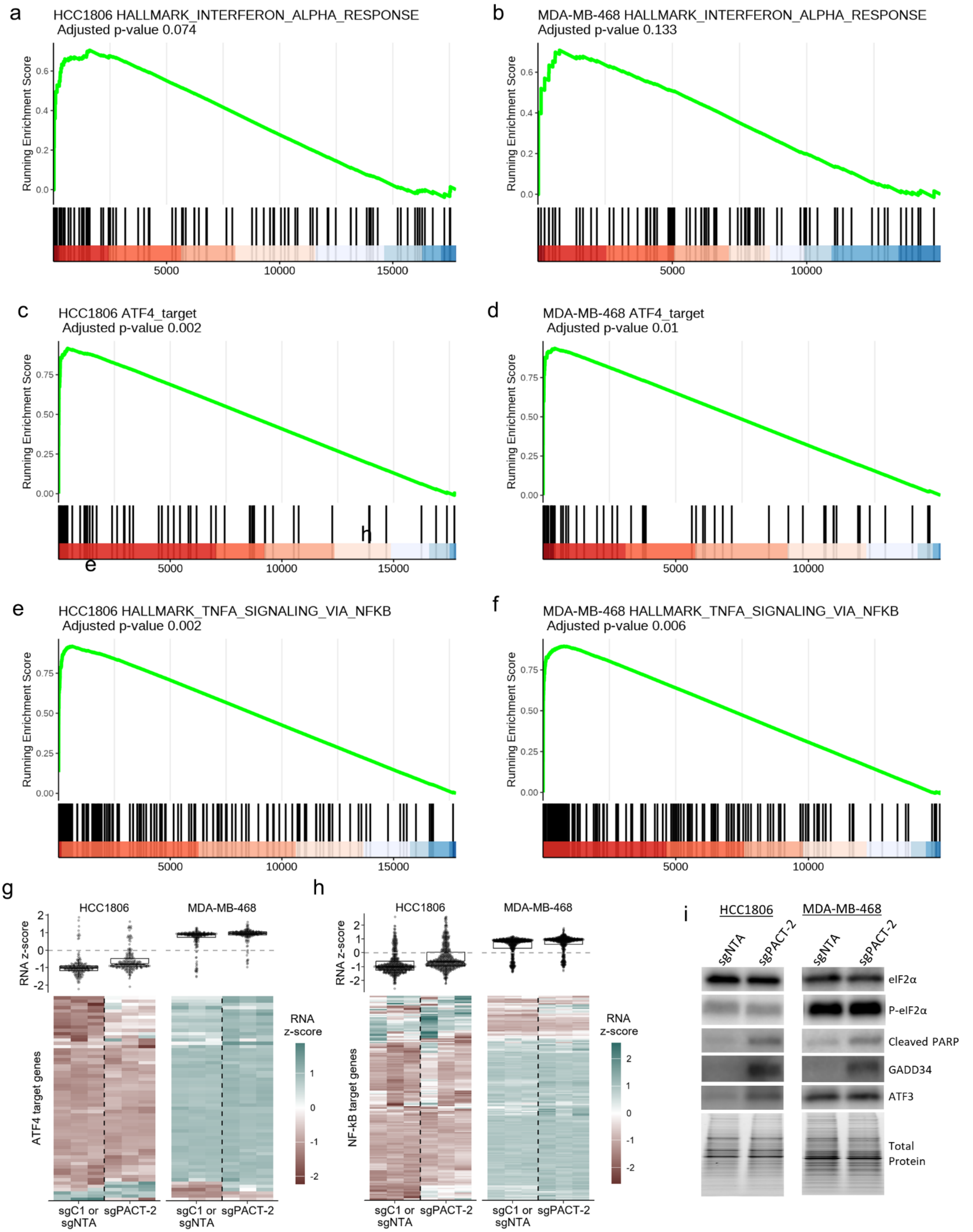
a
HCC1806 HALLMARK_INTERFERON_ALPHA_RESPONSE
Adjusted p-value 0.074
Running Enrichment Score
b
MDA-MB-468 HALLMARK_INTERFERON_ALPHA_RESPONSE
Adjusted p-value 0.133
c
HCC1806 ATF4_target
Adjusted p-value 0.002
d
MDA-MB-468 ATF4_target
Adjusted p-value 0.01
h
e
e
HCC1806 HALLMARK_TNFA_SIGNALING_VIA_NFKB
Adjusted p-value 0.002
f
MDA-MB-468 HALLMARK_TNFA_SIGNALING_VIA_NFKB
Adjusted p-value 0.006
5000
10000
15000
g
HCC1806
MDA-MB-468
RNA z-score
ATF4 target genes
RNA
z-score
sgC1 or sgNTA
sgPACT-2
h
NF-kB target genes
i
HCC1806
MDA-MB-468
sgNTA
sgPACT-2
eIF2α
P-eIF2α
Cleaved PARP
GADD34
ATF3
Total Protein

**Figure S3**

**a**-**f** GSEA plots for relevant gene sets. **g**-**h** Heatmaps for RNA expression of ATF4 or NF-κB target genes. Top panel, box and overlayed 'quasirandom' plots for all genes in the heatmap below. The heatmap is clustered by gene (rows), the dendrogram has been omitted for brevity. **i** Representative immunoblot of PACT depleted and control cells. The blots for each cell line were performed simultaneously, but exposure times for some proteins vary, as such bands should not be compared between cell lines. Total protein was imaged using a Stain-Free Gel and was used as the loading control for normalization.

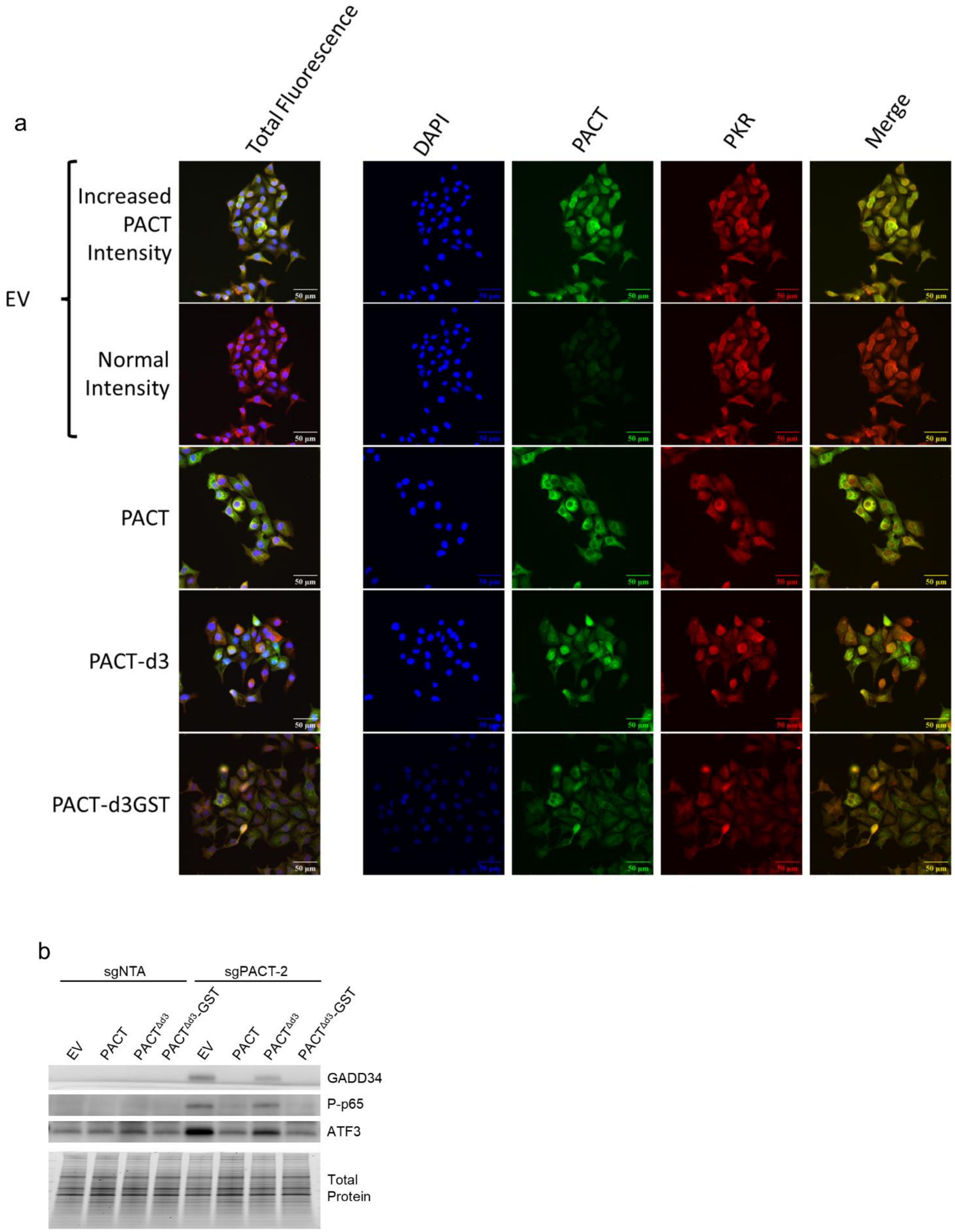


**Figure S4**

**a** Representative immunofluorescence micrographs for EV and PACT overexpression HCC1806 cells. For PACT, PACT-d3, PACT-d3GST, the signal acquisition parameters were the same as EV – Normal Intensity. **b** Representative immunoblot for control (sgNTA) and PACT depleted (sgPACT-2), with (PACT, PACT$^{\Delta d3}$ or

**Figure S4-cont**

PACT$^{\Delta d3}$-GST) or without (EV, empty vector) overexpression of WT or truncated PACT. Total protein is a Stain-Free gel image used as a loading control.

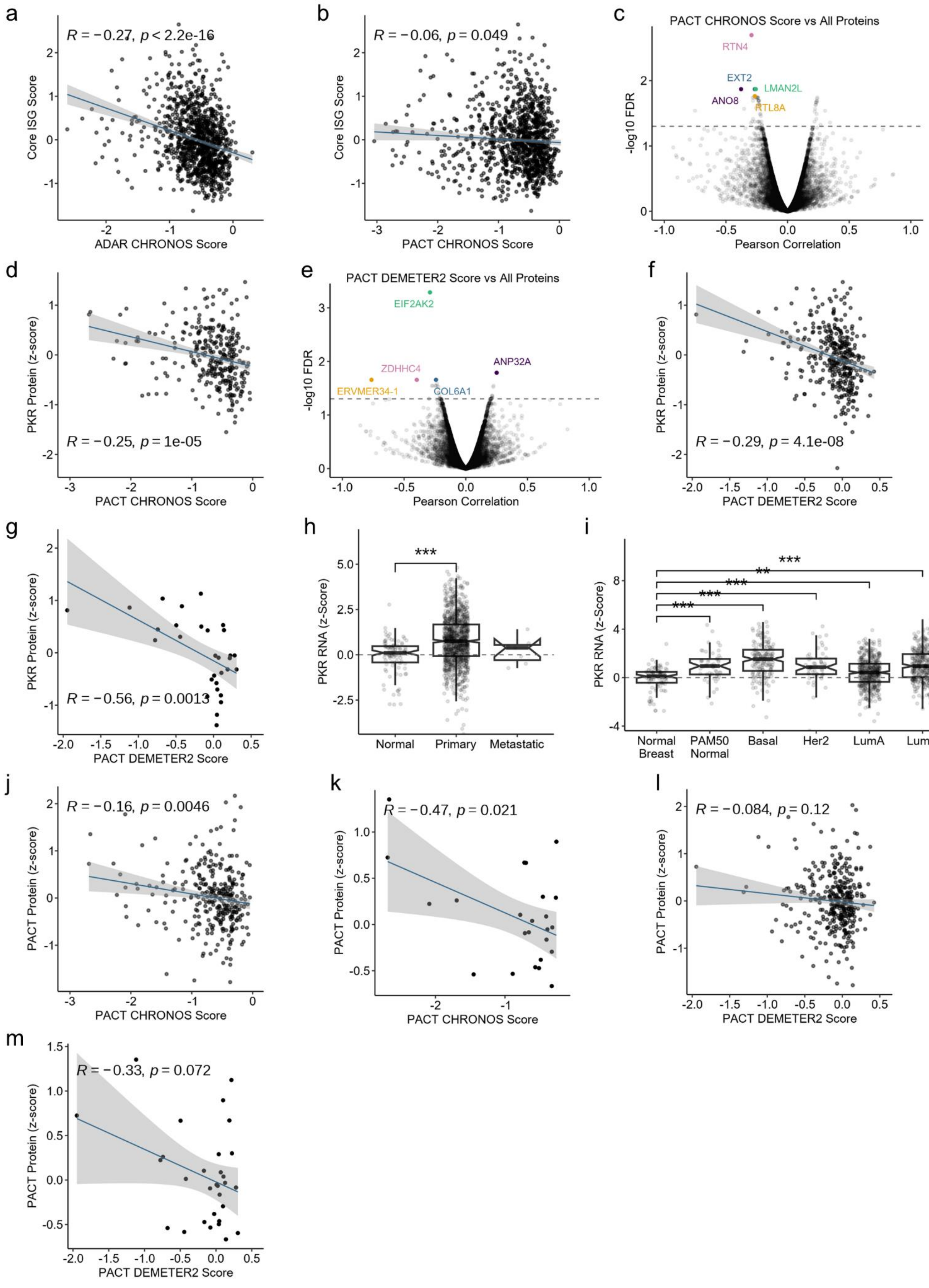
a
R = −0.27, p < 2.2e-16
Core ISG Score
ADAR CHRONOS Score
b
R = −0.06, p = 0.049
Core ISG Score
PACT CHRONOS Score
c
PACT CHRONOS Score vs All Proteins
RTN4
EXT2
LMAN2L
ANO8
RTL8A
-log10 FDR
Pearson Correlation
d
PKR Protein (z-score)
R = −0.25, p = 1e-05
PACT CHRONOS Score
e
PACT DEMETER2 Score vs All Proteins
EIF2AK2
ZDHHC4
ANP32A
ERVMER34-1
COL6A1
-log10 FDR
Pearson Correlation
f
PKR Protein (z-score)
R = −0.29, p = 4.1e-08
PACT DEMETER2 Score
g
PKR Protein (z-score)
R = −0.56, p = 0.0013
PACT DEMETER2 Score
h
***
PKR RNA (z-Score)
Normal
Primary
Metastatic
i
***
**
***
***
***
PKR RNA (z-Score)
Normal Breast
PAM50 Normal
Basal
Her2
LumA
LumB
j
R = −0.16, p = 0.0046
PACT Protein (z-score)
PACT CHRONOS Score
k
R = −0.47, p = 0.021
PACT Protein (z-score)
PACT CHRONOS Score
l
R = −0.084, p = 0.12
PACT Protein (z-score)
PACT DEMETER2 Score
m
R = −0.33, p = 0.072
PACT Protein (z-score)
PACT DEMETER2 Score

**Figure S5**

**Figure S5-cont.**

**a** – **b** Scatter plots comparing ISG expression (Core ISG Score, described previously (Kung et al. 2021)) and either ADAR1-dependency score (ADAR CHRONOS Score, **a**) or PACT-dependency score (PACT CHRONOS Score, **b**) for all cancer cell lines in DepMap. **c** and **e** Volcano plots of Pearson correlation coefficients and FDR corrected p-values for pairwise comparisons between PACT CHRONOS score (**c**) or PACT DEMETER2 score (**e**) and protein abundance for all proteins in DepMap across all cell lines. **d**, **f** and **g** Scatter plots comparing PKR protein abundance and PACT-dependency score for all cell lines in DepMap (**d** and **f**) or only breast cancer cell lines (**g**). **h** and **i** PKR RNA expression in normal human breast and breast tumors, data from TCGA. **j**-**m**, Scatter plots comparing PACT protein abundance and PACT-dependency score for all cell lines in DepMap (**j** and **l**) or only breast cancer cell lines (**k** and **m**). For all scatter plots, the Pearson correlation coefficient and p-values are shown. For panels **h**-**i**, data from TCGA; all other panels DepMap.

Below are uncropped blots for the main and supplemental figures. Some blots have multiple exposures. The exposure used for each protein in the main or supplemental figure is labeled in bold. Stain-free gel images for total protein are included and annotated to describe for which blots they correspond.

### FIGURE 2a/9e

Figure 2a

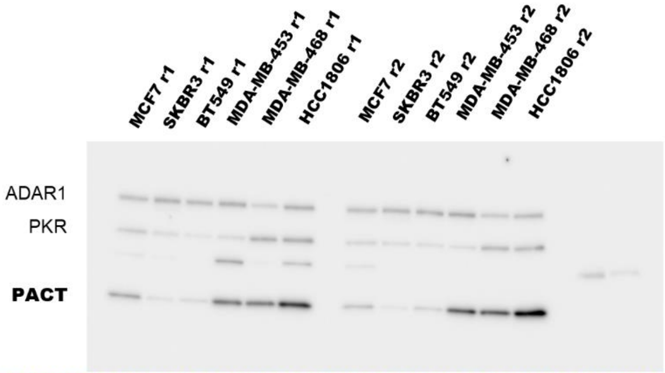


These two rightmost lanes were part of an unrelated experiment and not presented in the figure.

The immunoblot here is used for ADAR1 in 2a and PKR in 9e. Both were quantified using the gel below.

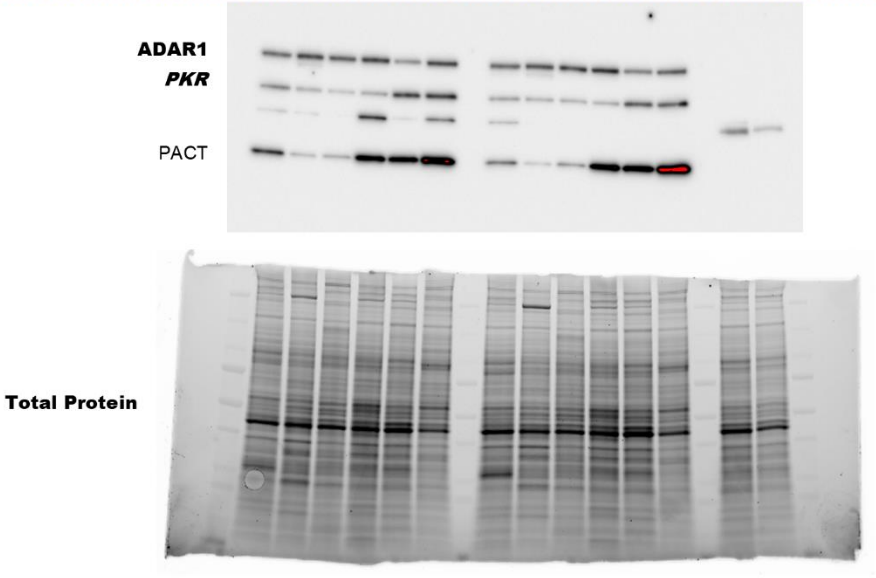


This gel is used for the normalization of ADAR1, PKR, and PACT.

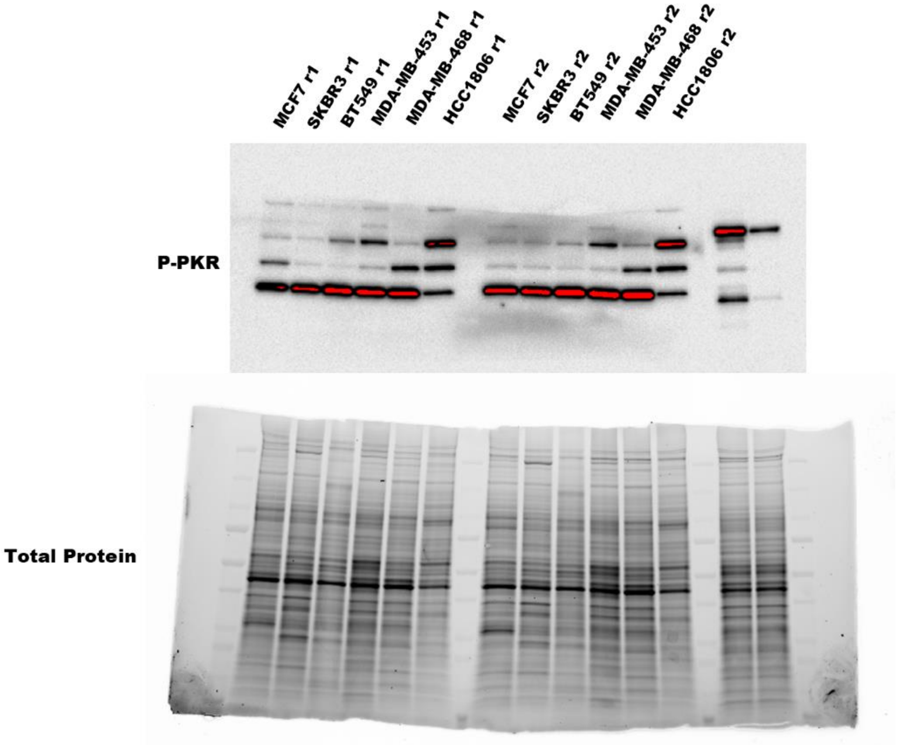


This gel is used for the normalization of P-PKR

Figure 9e

FIGURE 3a.

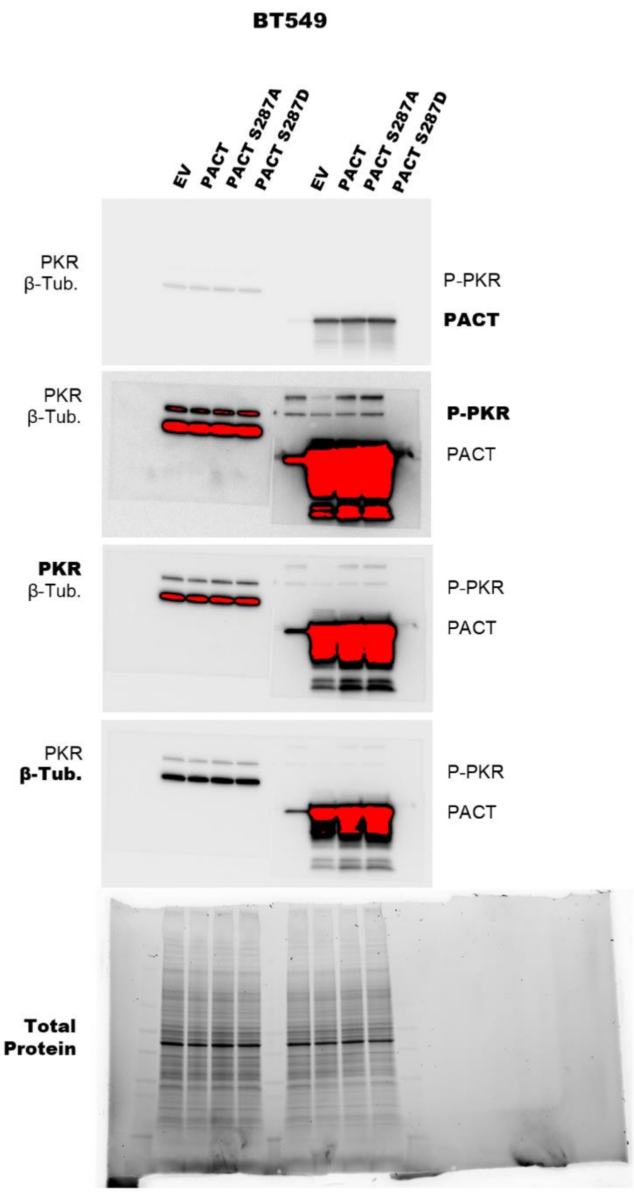
BT549
EV
PACT
PACT S287A
PACT S287D
EV
PACT
PACT S287A
PACT S287D
PKR
β-Tub.
P-PKR
PACT
PKR
β-Tub.
P-PKR
PACT
PKR
β-Tub.
P-PKR
PACT
PKR
β-Tub.
P-PKR
PACT
Total Protein

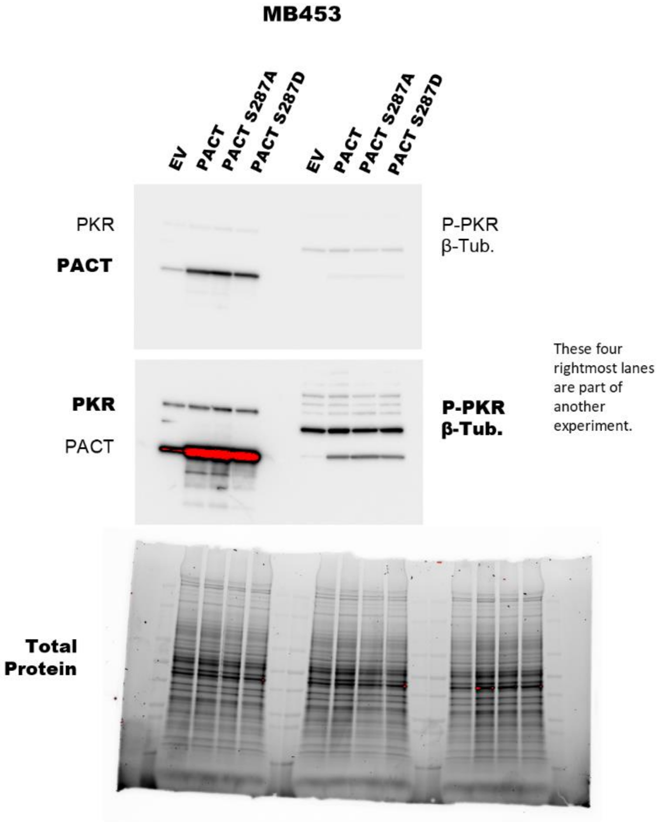
MB453
EV
PACT
PACT S287A
PACT S287D
EV
PACT
PACT S287A
PACT S287D
PKR
PACT
P-PKR
β-Tub.
PKR
PACT
P-PKR
β-Tub.
These four rightmost lanes are part of another experiment.
Total Protein

### FIGURE 3a-(dependent).

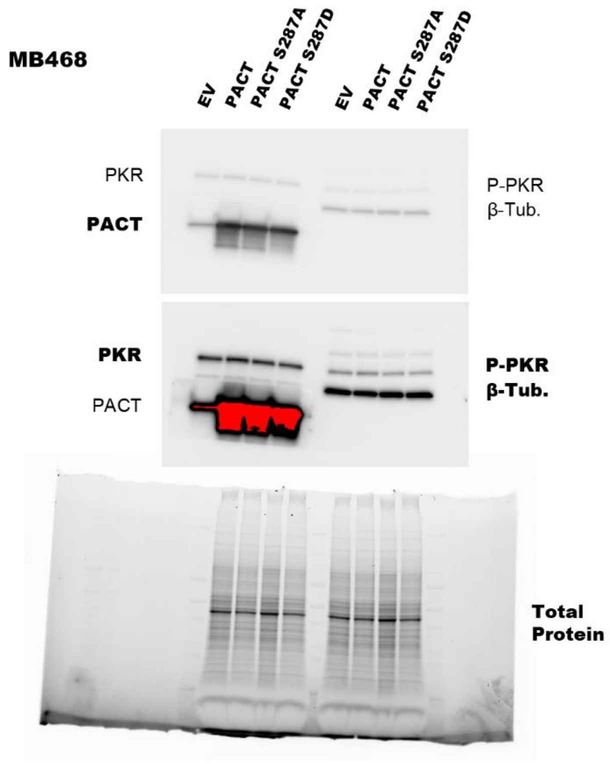


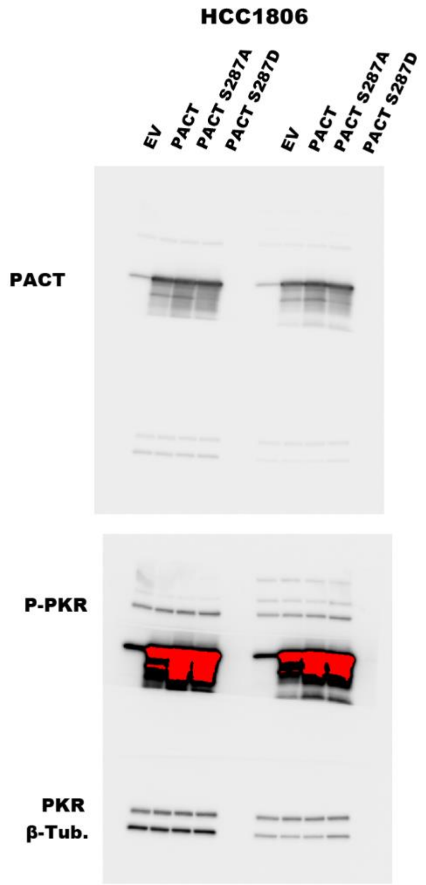


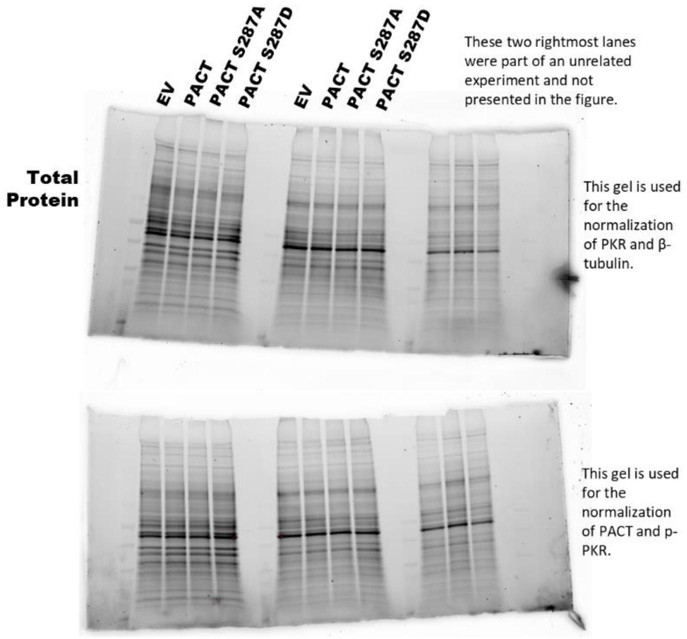

### FIGURE 4b-(independent).

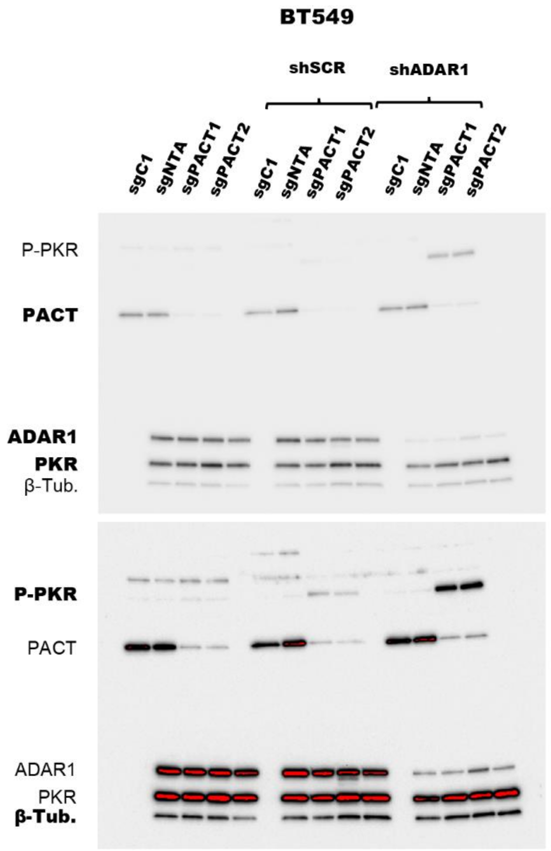


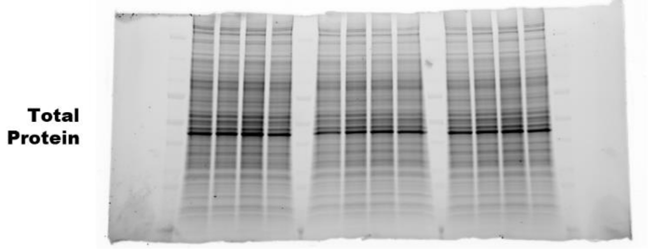


This gel is used for the normalization of P-PKR and PACT.

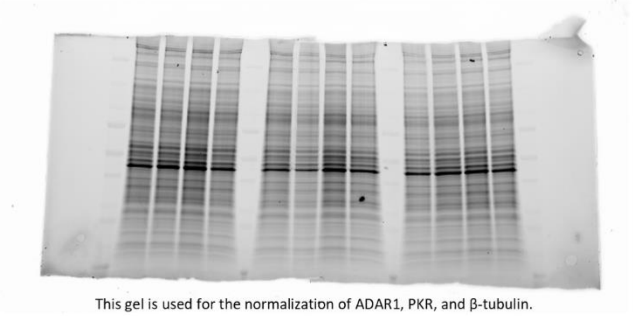

This gel is used for the normalization of ADAR1, PKR, and β-tubulin.

This gel and immunoblot is the same as that used in Figure 8a

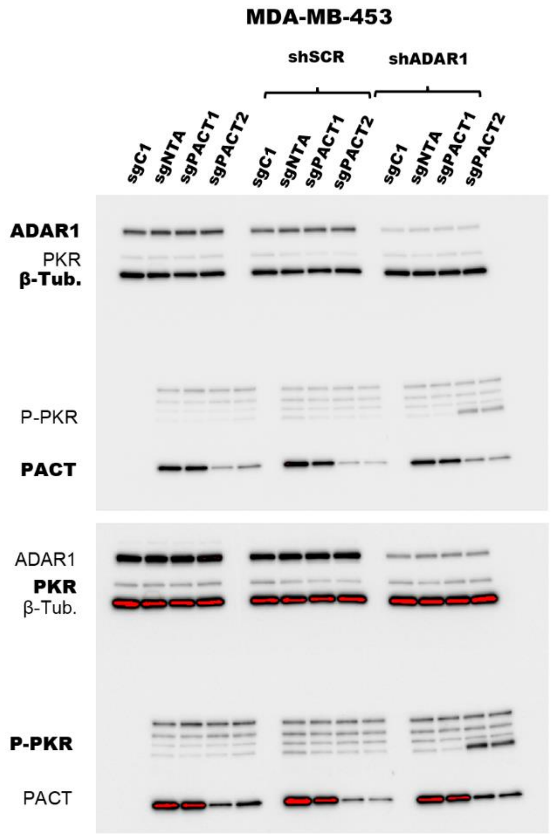


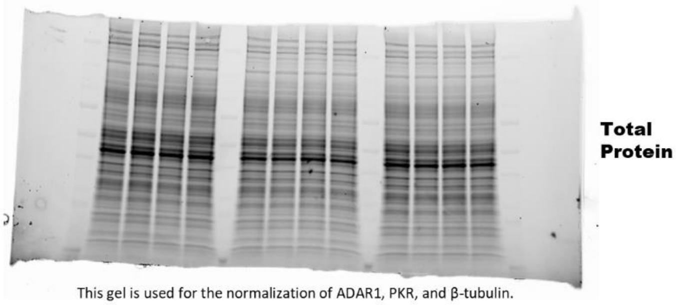


This gel is used for the normalization of ADAR1, PKR, and β-tubulin.

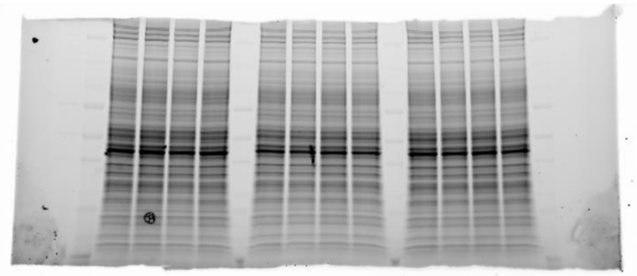

This gel is used for the normalization of P-PKR and PACT.

### FIGURE 4b-(dependent)

#### MDA-MB-468

The leftmost replicate is presented in the main figure.

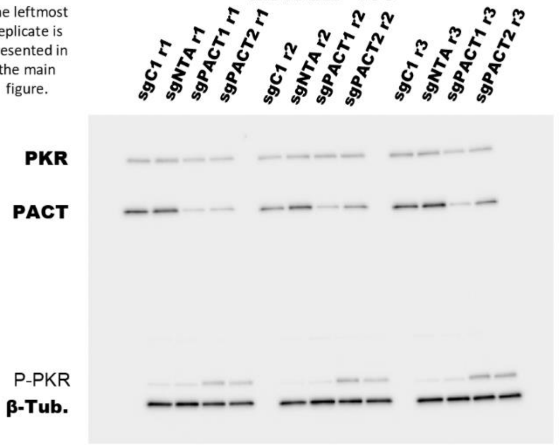


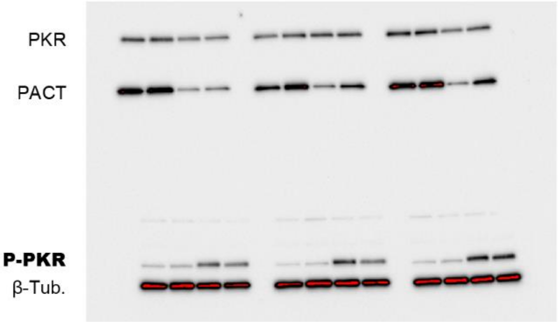


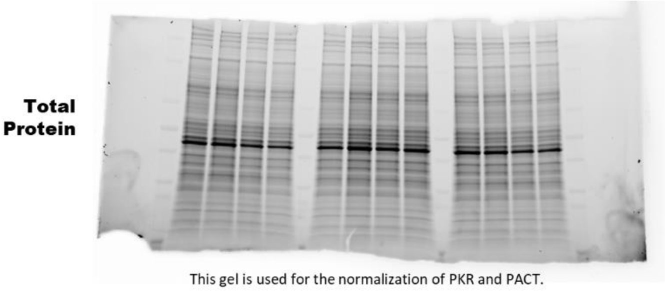


This gel is used for the normalization of PKR and PACT.

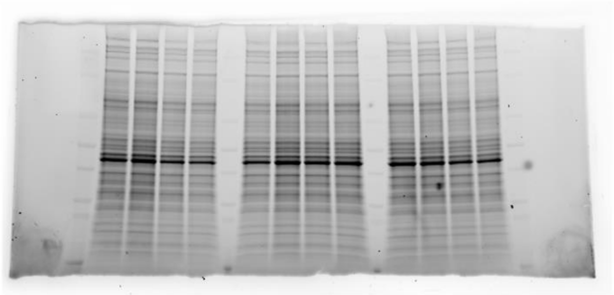

This gel is used for the normalization of P-PKR and β-tubulin.

#### HCC1806

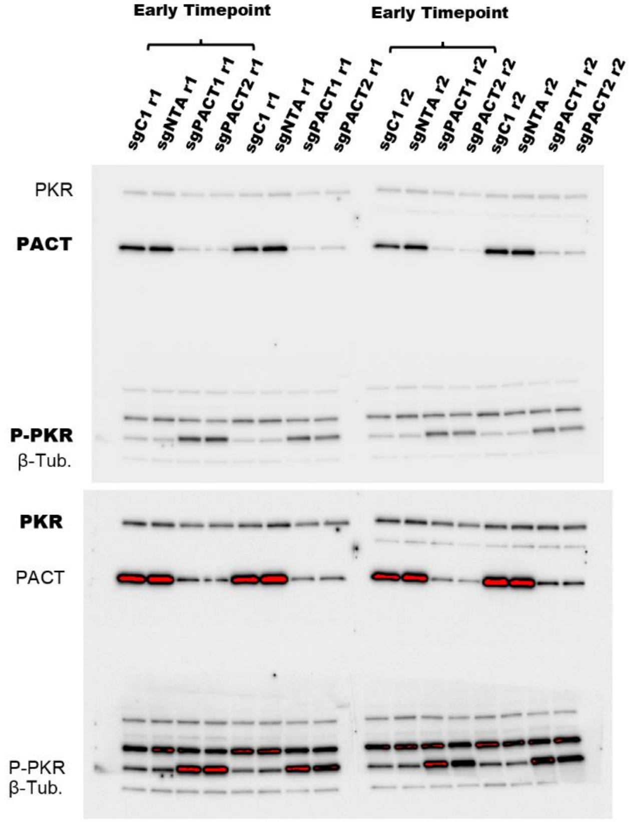


β-tubulin was re-probed and imaged a second time.

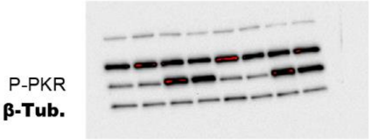


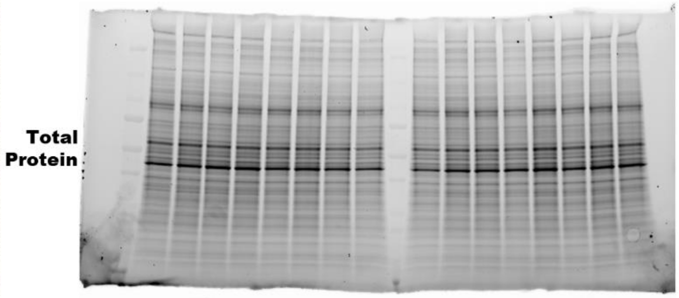


This gel is used for the normalization of P-PKR and β-tubulin.

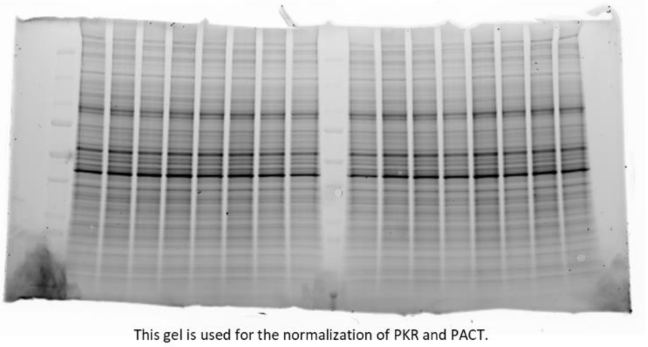

This gel is used for the normalization of PKR and PACT.

### FIGURE 6d.

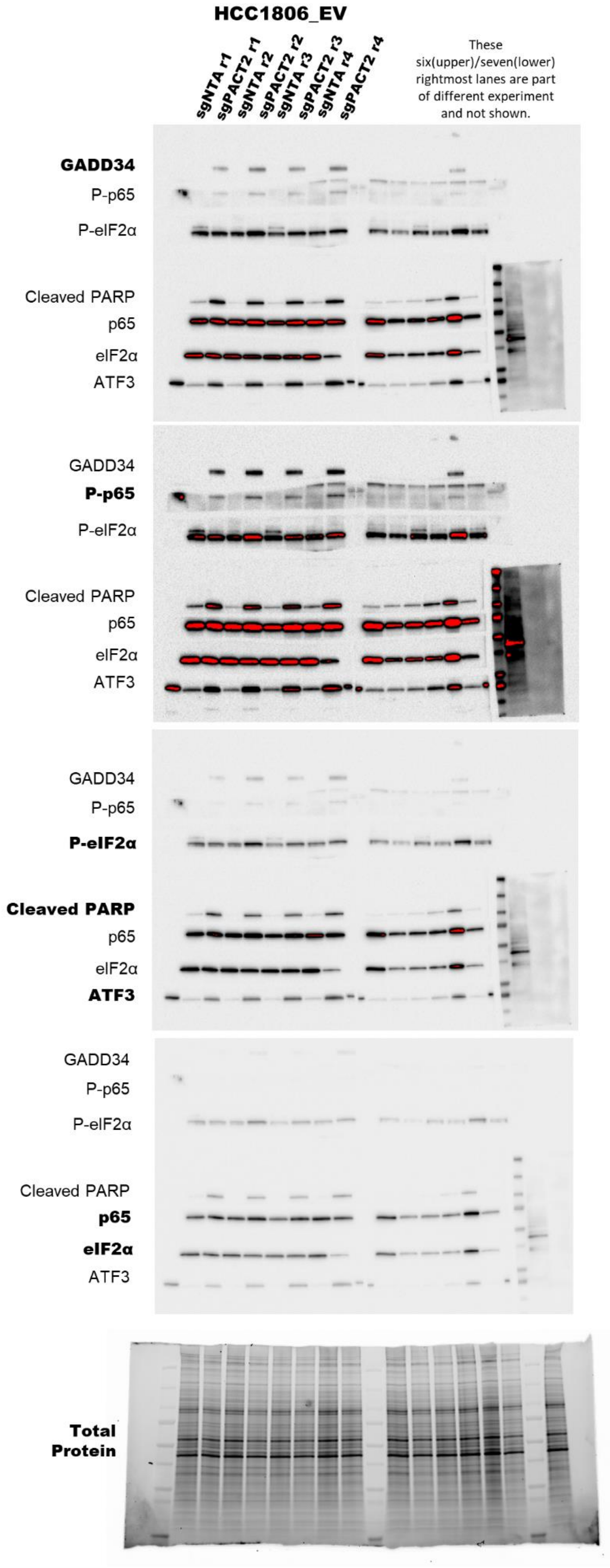


This gel is used for the normalization of Cleaved PARP, p65, eIF2α, and ATF3.

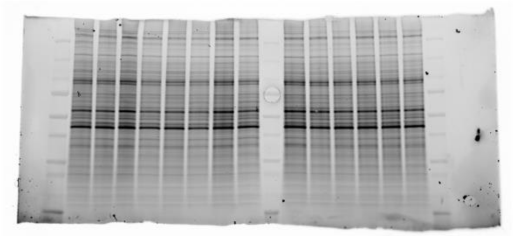

This gel is used for the normalization of GADD34, P-p65, and P-eIF2α.

### FIGURE S3i.

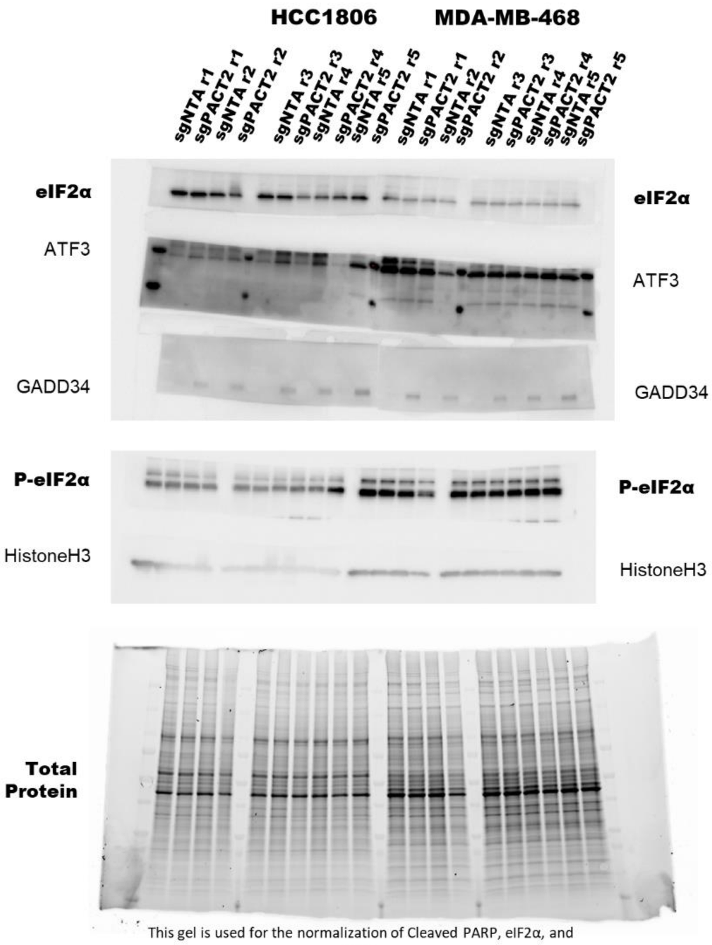


This gel is used for the normalization of Cleaved PARP, eIF2α, and ATF3.

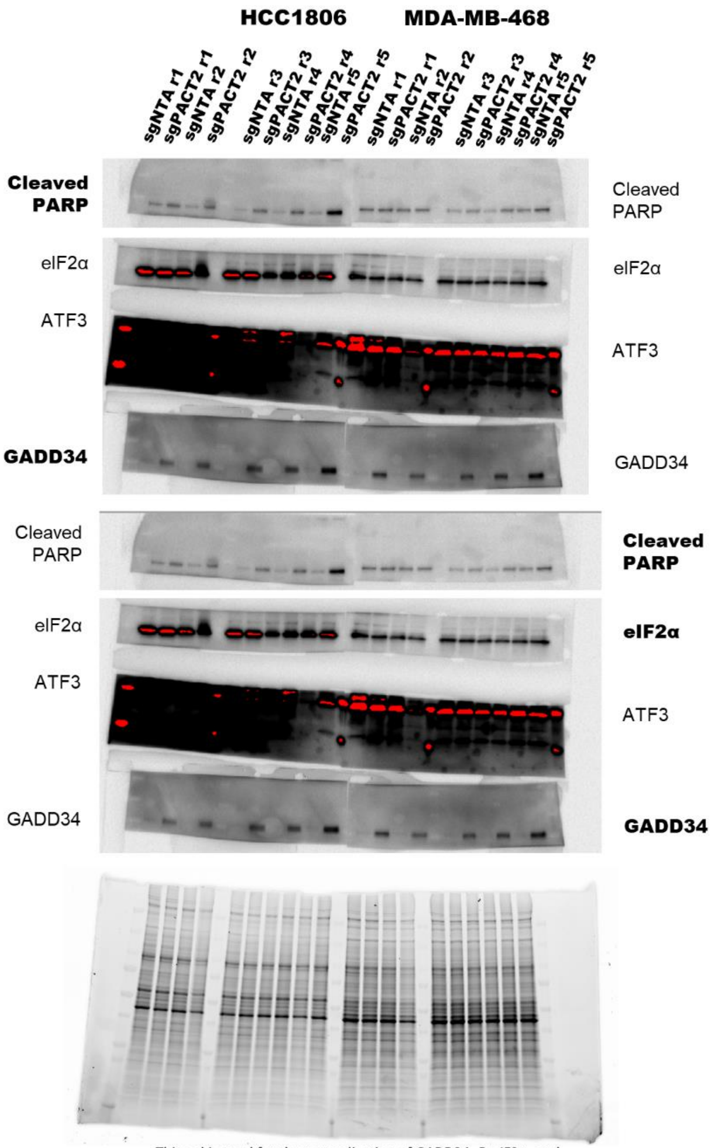


This gel is used for the normalization of GADD34, P-eIF2α, and HistoneH3.

FIGURE 6e.

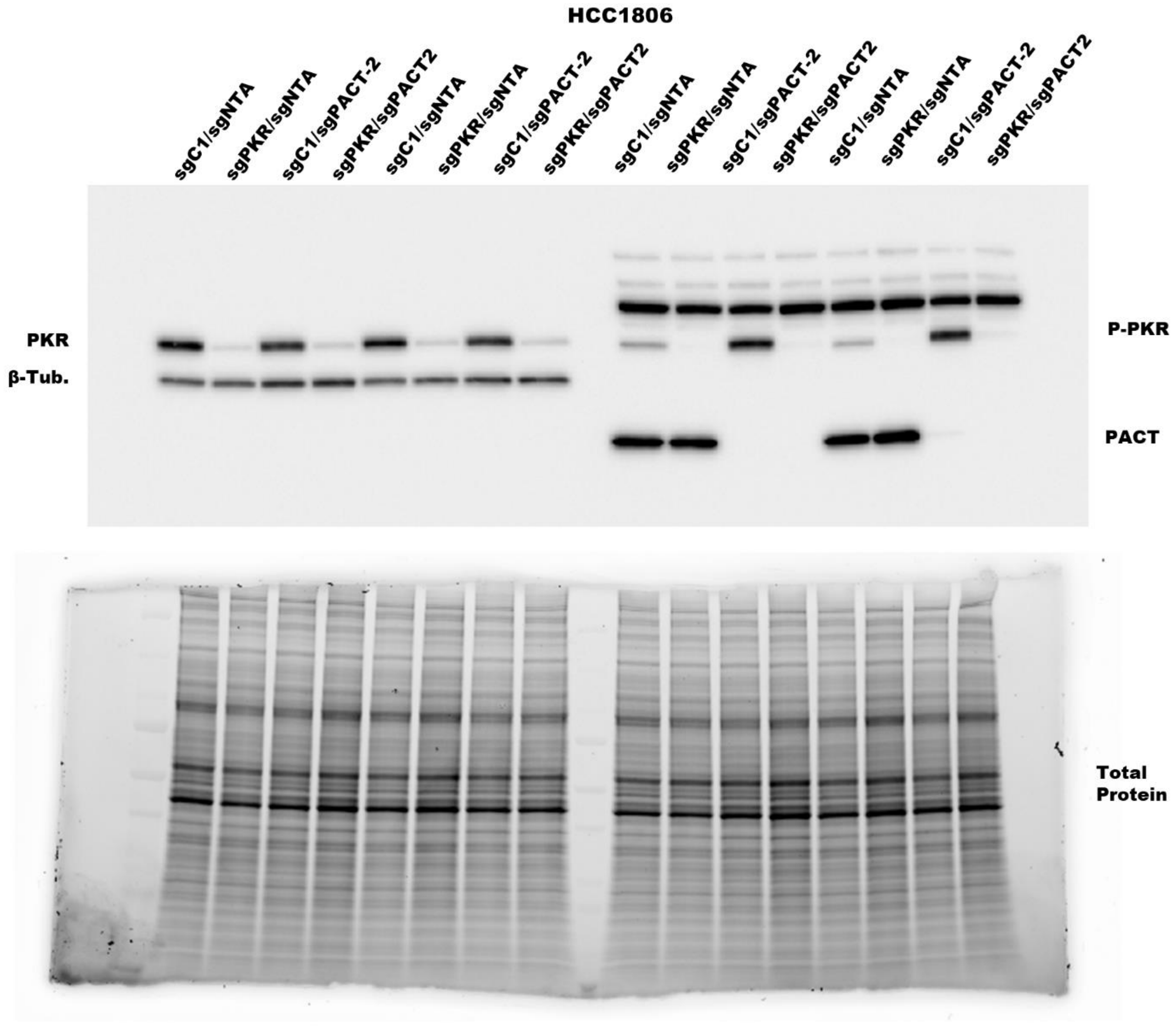
HCC1806
sgC1/sgNTA
sgPKR/sgNTA
sgC1/sgPACT-2
sgPKR/sgPACT2
sgC1/sgNTA
sgPKR/sgNTA
sgC1/sgPACT-2
sgPKR/sgPACT2
sgC1/sgNTA
sgPKR/sgNTA
sgC1/sgPACT-2
sgPKR/sgPACT2
sgC1/sgNTA
sgPKR/sgNTA
sgC1/sgPACT-2
sgPKR/sgPACT2
PKR
β-Tub.
P-PKR
PACT
Total
Protein

FIGURE 7b.

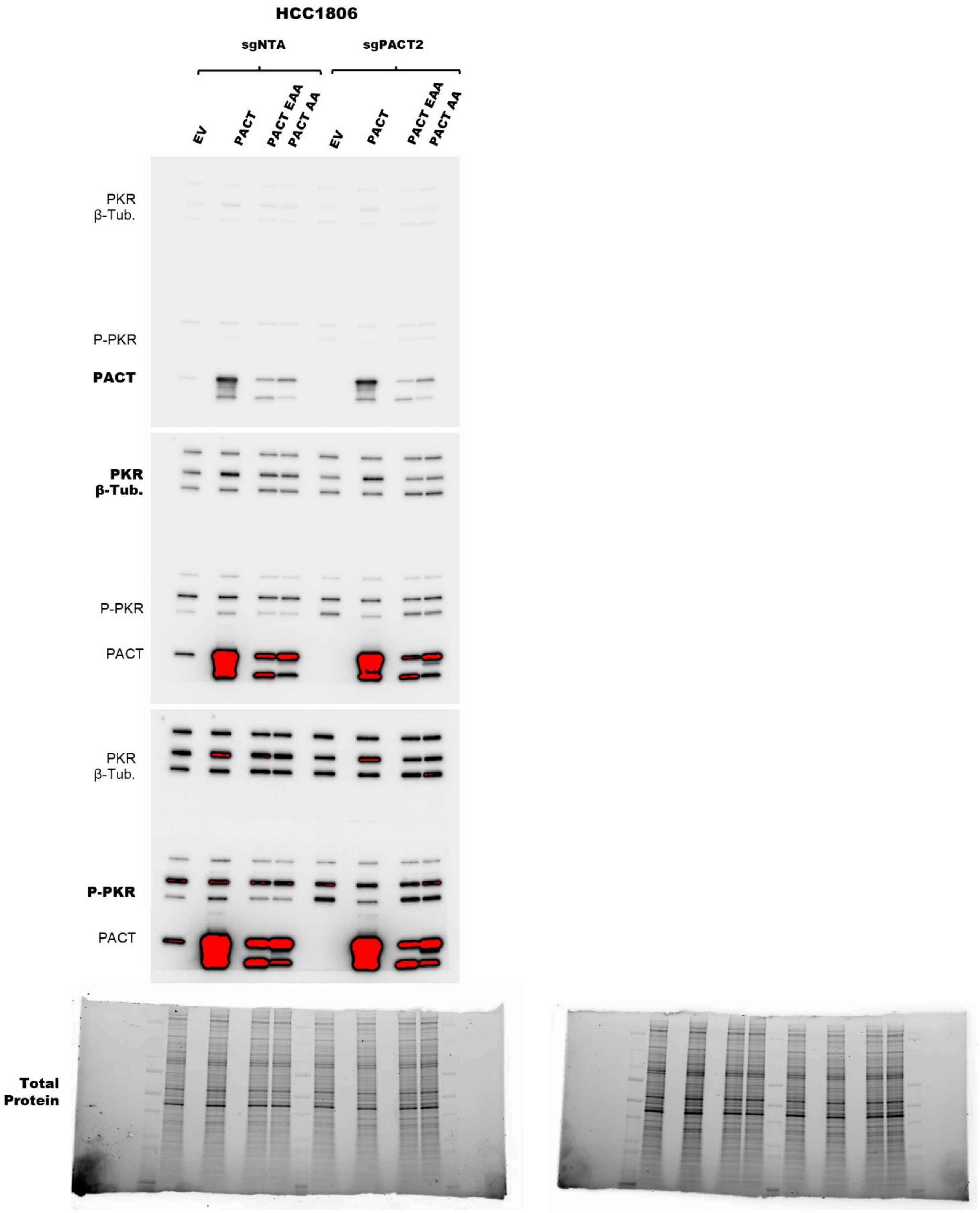


This gel is used for the normalization of PKR and β-tubulin.

This gel is used for the normalization of P-PKR and PACT.

### FIGURE 7f.

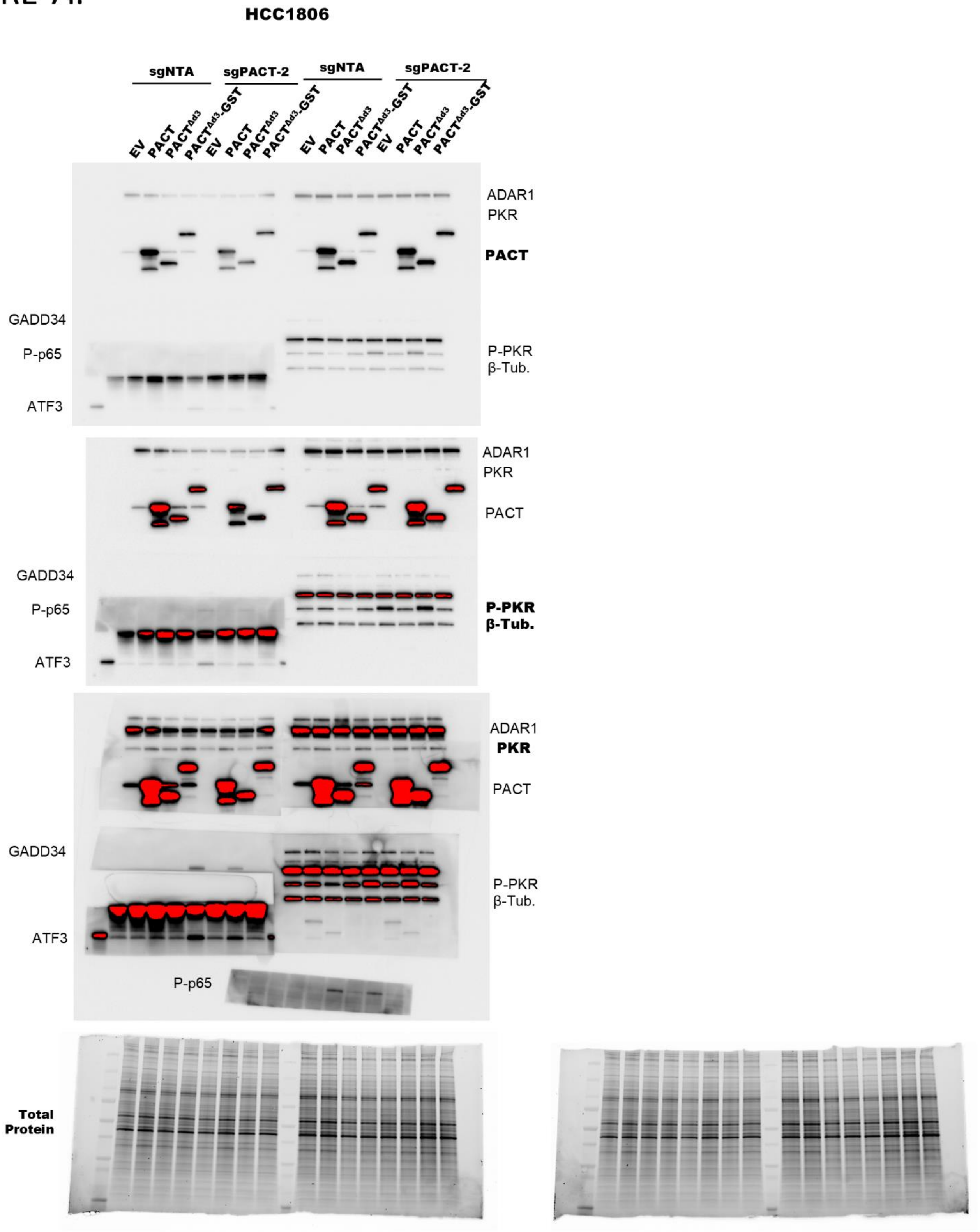


This gel is used for the normalization GADD34, P-p65, ATF3, P-PKR, and β-tubulin.

This gel is used for the normalization of ADAR, PKR, and PACT.

### FIGURE 8a.

This gel and immunoblot is the same as that used in Figure 4b

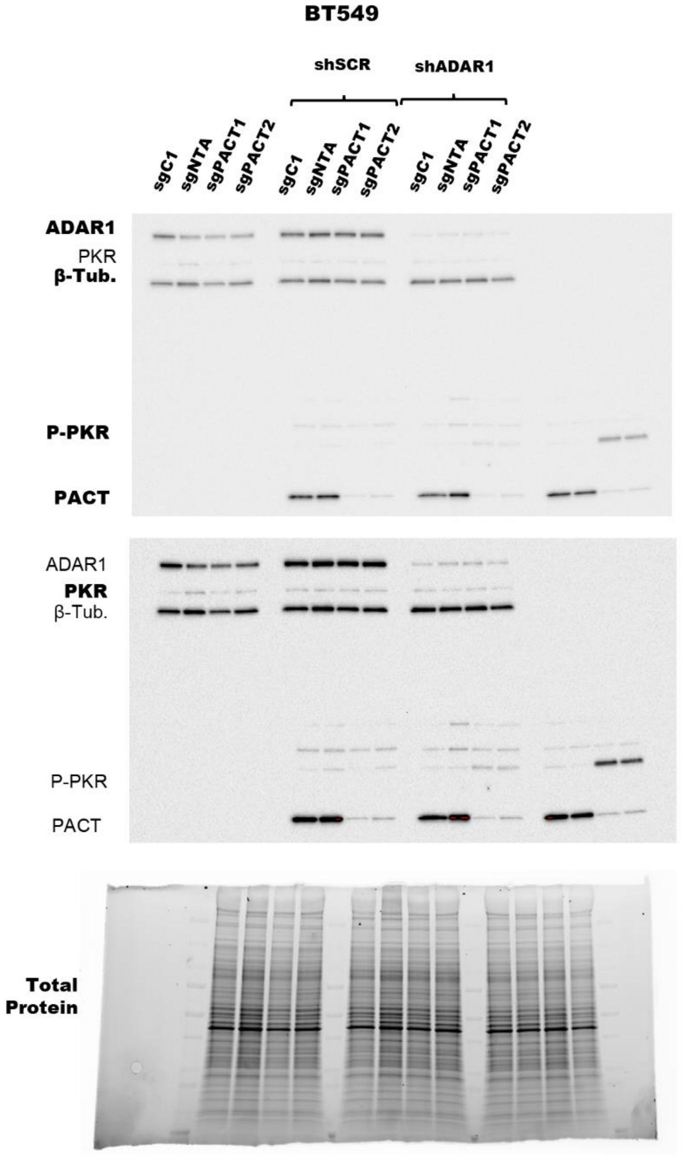


This gel is used for the normalization of ADAR1, PKR, and β-tubulin.

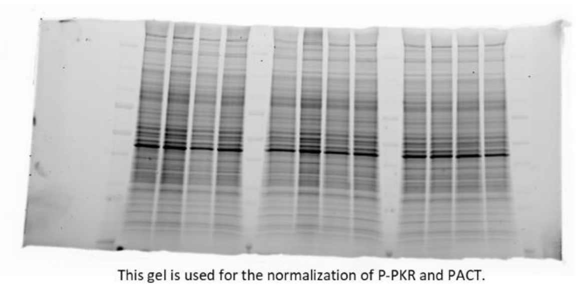

This gel is used for the normalization of P-PKR and PACT.

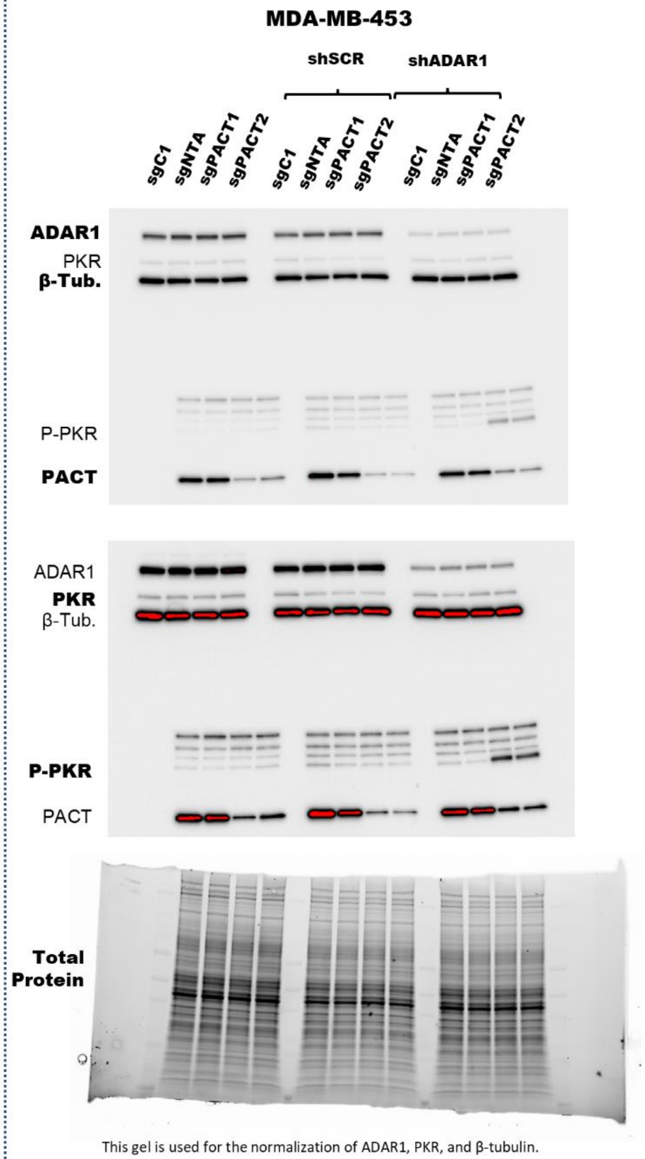


This gel is used for the normalization of ADAR1, PKR, and β-tubulin.

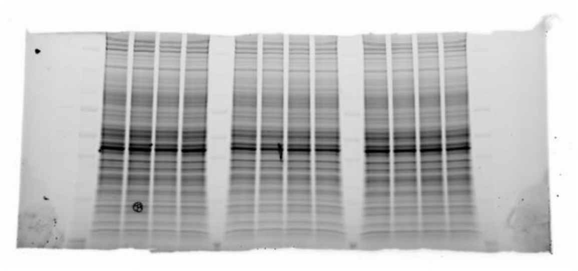

This gel is used for the normalization of P-PKR and PACT.

### FIGURE 8h.

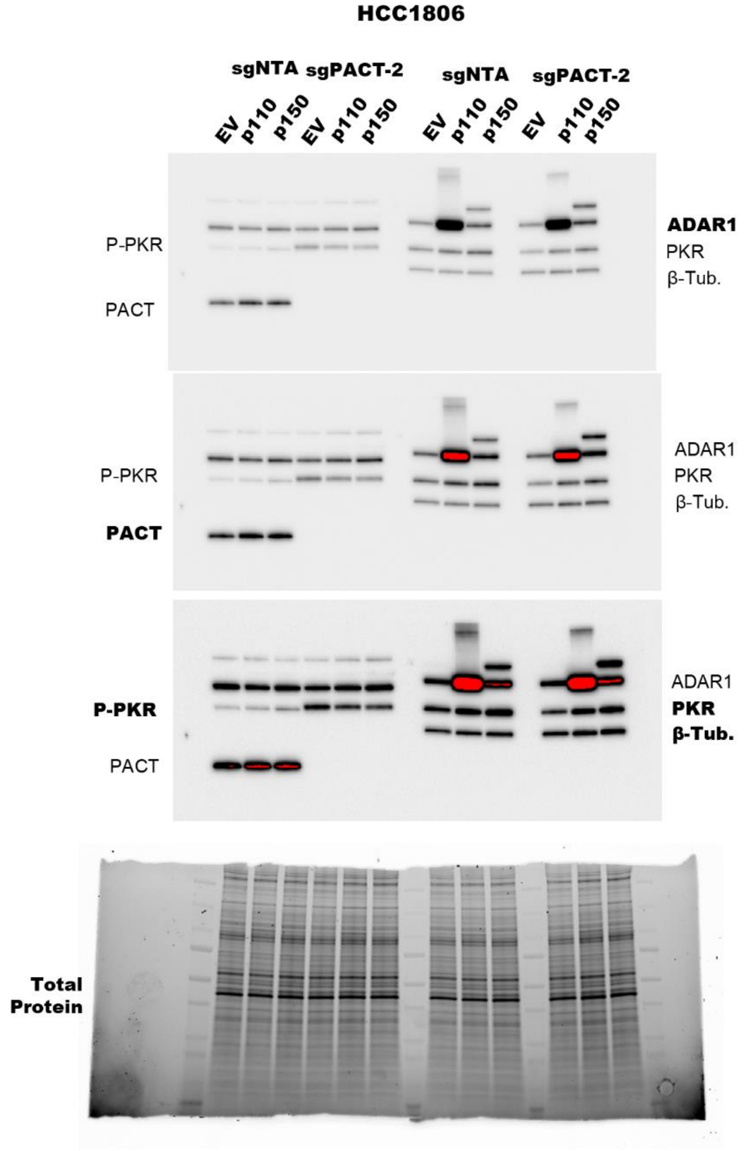

HCC1806
sgNTA
sgPACT-2
sgNTA
sgPACT-2
EV
p110
p150
EV
p110
p150
EV
p110
p150
EV
p110
p150
P-PKR
PACT
ADAR1
PKR
β-Tub.
P-PKR
PACT
ADAR1
PKR
β-Tub.
P-PKR
PACT
ADAR1
PKR
β-Tub.
Total Protein
